# Binding of different substrate molecules at the docking site and the active site of γ-secretase can trigger toxic events in sporadic and familial Alzheimer’s disease

**DOI:** 10.1101/2022.04.12.487996

**Authors:** Željko M. Svedružić, Vesna Šendula Jengić, Lucija Ostojić

## Abstract

**Significance:** pathogenic changes in γ-secretase activity, and its response to different drugs, can be greatly affected by changes in saturation of γ-secretase with its substrate. The molecular mechanism is unclear.

**Results:** multiscale molecular dynamics studies show that saturation of γ-secretase with its substrate can result in parallel binding of different substrate molecules at the docking site and the active site. C-terminal domain of the second substrate can bind at cytosolic end of presenilin subunit while γ-secretase is still processing its first substrate. Such interactions can disrupt dynamic presenilin structures that regulate proteolytic steps. Similar disruptions in dynamic presenilin structures can be produced by different drugs and by different disease-causing mutations. Thus, the presented two-substrate mechanism, can explain toxic inhibition of γ-secretase activity and toxic increase in production of the longer, more hydrophobic, Aβ-proteins. Toxic aggregation between N-terminal domains of the two substrates is controlled by nicastrin ectodomain. Such aggregation is more likely to happen with C99-βCTF-APP than with C83-αCTF-APP substrate, which can explain, why β-secretase path is more pathogenic than α-secretase path. The binding of C99-βCTF-APP substrate to γ-secretase can be controlled by substrate-channeling between nicastrin ectodomain and β-secretase.

**Conclusions:** The presented two-substrate mechanism can explain why different studies consistently show that increase in saturation of γ-secretase with its substrate can support pathogenic changes in different sporadic and familiar cases of the disease. Future drug-development strategies can target different physiological mechanisms that control the balance between cellular levels of γ-secretase activity and the total amyloid metabolism.

## Introduction

Alzheimer’s disease is slowly progressing and ultimately fatal neurodegenerative disorder [1,2]. Alzheimer’s disease stands out ahead of malignant diseases, as the biggest financial burden for health care providers in developed countries [1,3,4]. Impressive drug-development efforts have been mostly centered on the metabolism of the last 99 amino acids of amyloid precursor protein (C99-βCTF-APP) [3,4]. The most frequent targets are two aspartic proteases: membrane-anchored β-secretase, and membrane-embedded γ-secretase [2,3,5]. A number of compounds have been developed, with different structures, different binding sites, different mechanism of action, different pharmacological properties, and very impressive nanomolar potency [1,4,5]. The impressive list of diverse and potent compounds did not give desired results, but it clearly showed, that the present challenges must go beyond routine medicinal chemistry. It appears that β-secretase and γ-secretase have some specific features in the enzymatic mechanism, which represent some unique challenges for the drug-design efforts [4,6-14].

Several pathogenic changes in Aβ production can be observed when γ-secretase is gradually saturated with its substrate [7,10,13,14]. Changes in saturation of γ-secretase with its substrate can also significantly affect how the enzyme responds to potential drugs [6,9,11,12]. The earliest age-of-onset can be observed with mutants that have the biggest chance to reach saturation at the lowest substrate loads [7]. Changes in saturation of γ-secretase with its substrate is a key physiological process [15], sadly the underlying mechanism is not clear.

Different studies of γ-secretase activity have never been satisfactorily explained [16], because mechanistic interpretations of enzyme activity studies depend on quantitative analysis [9,11-13]. Quantitative analysis of complex enzyme activity depends on mathematical modeling [17-19]. A significant fraction of biomedical scientists is not fully familiar with such methods [17-19]. Fortunately, computational studies of molecular structures, can greatly simplify and advance interpretation of the enzyme activity and drug-design studies [8,20-26]. In this study, we expand some of the earlier enzyme activity studies using advanced computational methods for the analysis of the interaction between γ-secretase, β-secretase, and C99-βCTF-APP [27,28]. Specifically, we analyze proposals that pathogenic events can be attributed to the accumulation of C99-βCTF-APP and Aβ molecules of different lengths [10,11,13,29].

γ-Secretase is a unique enzyme that has a separate substrate docking site and the active site [16,26,30]. In this study, we show that γ-secretase can bind in parallel different substrate molecules, at the docking site and the active site. The two substrates mechanism can happen when γ-secretase is exposed to an excess of its substrates, i.e. when substrate starts accumulating while the enzyme is still processing its substrate [9,10,17]. We show that the second substrate, different drugs, and different FAD mutations, all target the same sites on the presenilin subunit [8,31,32]. Thus, the binding of the second substrate, like different drugs and FAD mutations, can support the conformational changes that interfere with the processive catalysis and Aβ production [8,20,25,32]. We show that the presented two-substrate mechanism is consistent with different studies of changes in γ-secretase activity in Alzheimer’s disease. Building successful correlations between clinical and biochemical studies of Alzheimer’s disease is crucial for the development of effective early diagnostic methods and drug-development strategies [4].

## Results

### Multiscale molecular dynamics (MD) studies of dimerization of C99-βCTF-APP molecules in cholesterol-lipid-bilayer

Different structural studies showed that C99-βCTF-APP molecules are highly dynamic, readily affected by the experimental conditions, and difficult to measure [3,24,33-37]. Thus, we had to use multiscale MD studies, in combination with the earlier structural studies to describe possible interactions between C99-βCTF-APP molecules in a cholesterol-lipid-bilayer (Figs. 1 and 2, supp. video 1) [28].

**Figure 1.**
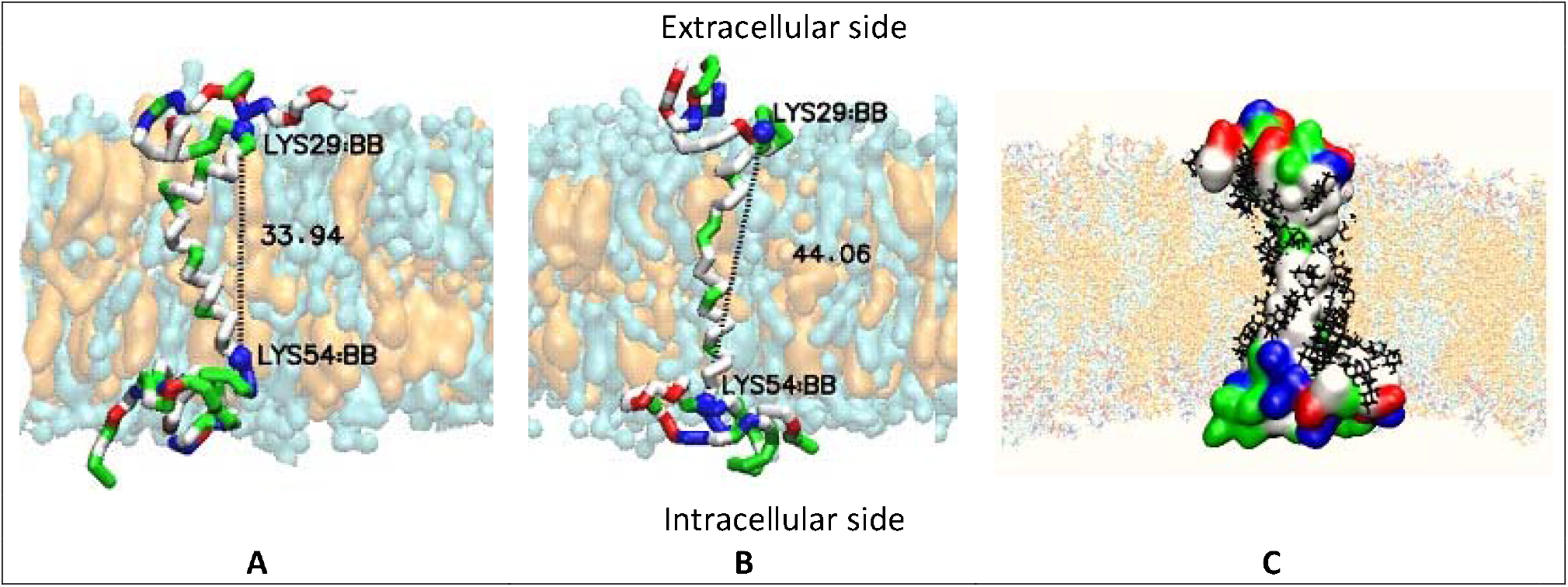
(<u>A</u>-<u>C</u>). Multiscale molecular dynamic studies of C99-β-CTF-APP structure in a cholesterol-lipid bilayer. Multiscale MD calculations can provide dynamic structural depictions of C99-β-CTF-APP structure that can be related to the earlier studies [24,33,37,100,101]. The amino acids are shown as hydrophobic (white), positive (blue), negative (red), and polar not charged (green). The cholesterol-lipid-bilayer shows surface models of cholesterol (orange) in a mixtur of POPC, POPA, POPE, POPS, POPI, and PSM molecules (cyan) (see methods). (**<u>A</u>**-**<u>B</u>**) The transmembrane section of the C99-β-CTF-APP backbone can exist in compact and extended forms [24]. The transmembrane helix is hydrophobic (white) with notable polar sites at Thr 43 and Thr 48 (green), and hinge sites at Gly38 and Gly39 (green). The extracellular and intracellular parts are rich in positive (blue), negative (red), and polar (green) amino acids. Changes in Lys29-Lys54 distances (numeration as in PDB:2LP1), showed that the shortest conformers are around 33.94 Å long, while the longest conformer is about 44.06 Å long. About 62% of the time protein takes conformations that are about 37.7 ± 2.5 Å long. (**<u>C</u>**) C99-β-CTF-APP surface shows that charged and polar amino acids form highly dynamic but compact structures in the extracellular and intracellular domains [24]. The transmembrane section can be covered with cholesterol molecules (black lines) [33].

**Figure 2.**
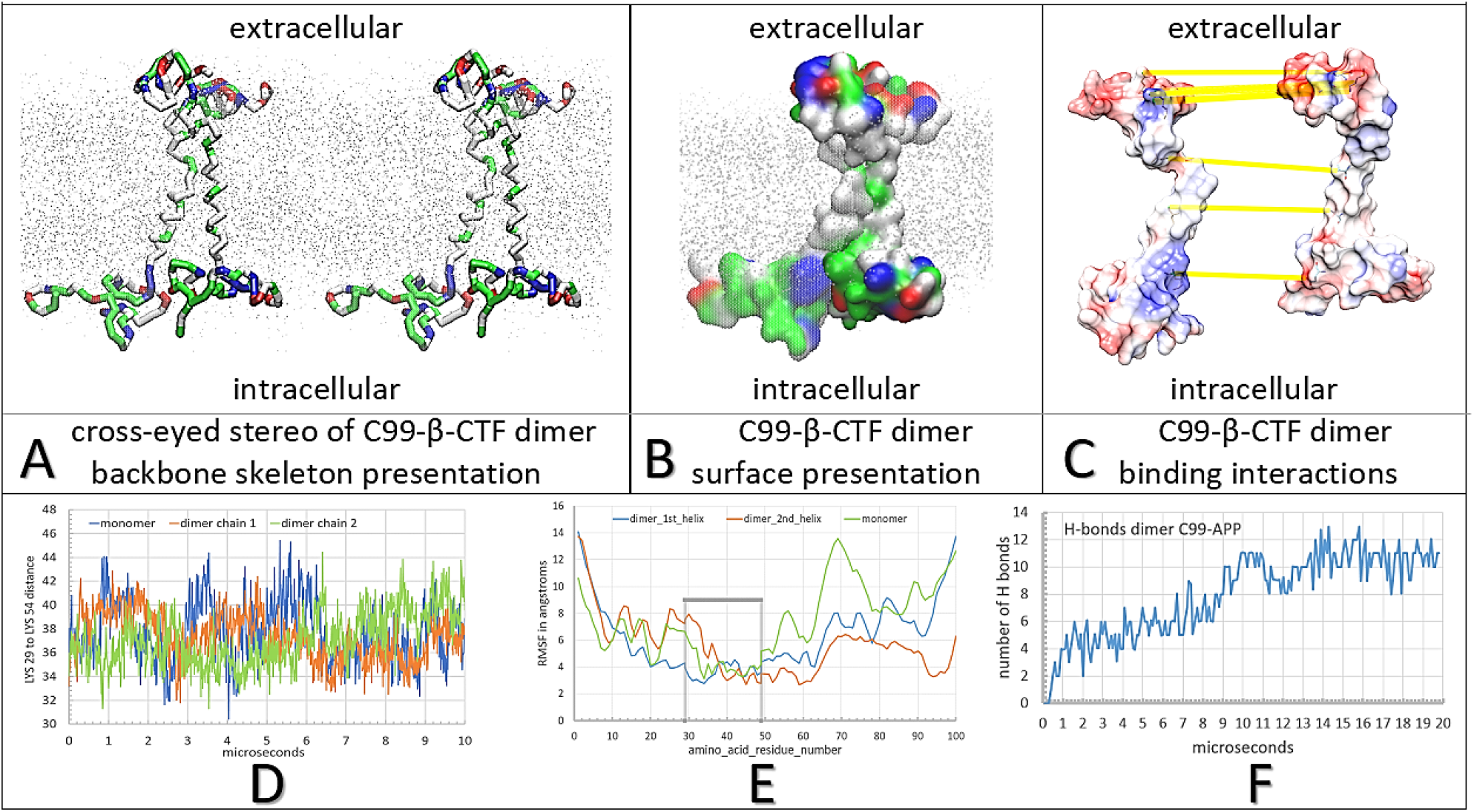
(<u>A</u>-<u>F</u>). Multiscale molecular dynamic studies of C99-β-CTF-APP dimerization in a cholesterol-lipid bilayer. The backbone models are used to show protein conformations, while the partially transparent Connolly surfaces are used to show protein-protein contacts [102]. The amino acids are colored as hydrophobic (white), positive (blue), negative (red), and polar not charged (green). For orientation Thr43 and Thr48 sites are visible as green sites in the TM section. Cholesterol-lipid-bilayer i shown as grey dots (methods). (**<u>A</u>**) cross-eyed stereo view of protein backbone can give the best insight into conformational changes that support formation of C99-β-CTF-APP dimer. Dimerization is initially driven by conformational changes at Gly38 and Gly39 sites (green), which act as hinge points. The final structure of the dimer is a result of multiple interactions between charged and polar amino acids at the C terminal and the N terminal domain. (**<u>B</u>**) surface models show that the two C99-β-CTF-APP molecules can form tight complementary surfaces down the full protein length. (**<u>C</u>**) the molecules in the complex are spread apart to show the analysis of complementary surfaces shapes and electric potentials (red-blue,-4.0 to 4.0 kBT/e). Highlighted are H bonds (yellow lines) [103]. The interaction sites on the C-terminal domain between the positive Lys53-Lys54-Lys55 site and negative site Glu74-Glu75 site. On the N-terminal domain highlighted are interactions sites between Glu4 and Lys16, and between Arg5 and Glu22-Asp23. (**<u>D</u>**) the changes in Lys29-Lys54 (numbering as in PDB: 2LP1) distances as a function of MD calculation steps showed that dimerization depends on combined contribution from the extended and compact forms of the substrate [24]. (**<u>E</u>**) RMSF values as a function of amino acid position show that dimerization is strongly dependent on the conformers in the polar parts of the substrate [39]. (**<u>F</u>**) changes in a number of H-bonds as a function of the calculated MD time. The initial lag represents initial diffusion and the first contact. The steps in the graph correspond to different conformers and local energy minima as two structures form the most stable complex.

We started all MD studies by building a full-length C99-βCTF-APP structure from the available NMR conformers (PDB: 2LP1, [33], see methods). With isolated C99-βCTF-APP molecule Lys29-Lys54 distances showed that the TM section constantly fluctuates between the two main conformations (Fig 1 A-B, numeration based on PDB: 2LP1, [33]). Similar to the earlier observations [24], we find that the TM sections can vary between: 45.45 Å long fully extended structure (Fig 1 B), and the shortest 33.09 Å long structure (Fig. 1A). Average distance is 37.7 ± 2.5 Å (Fig. 2 D). The compact forms are slightly more dominant, and occupy about 62% of the time. The conformers are mostly driven by the unusual hinge in the structure in the position of Gly37Gly38Val39Val40 sequence (Fig 1A, [24,33]). Ramachandran plots showed that the TM section has an α-helix structure (Fig 1 and 2 A-C). The OH groups on Thr 43 and Thr 48 are trapped in the hydrophobic environment (Fig 1 and 2 A-C), and must form hydrogen bonds with the adjacent peptide bond. We also found that the TM section gets readily covered on its surface by cholesterol as indicated earlier (Fig 1C, [37]).

MD studies show that initially extended structures in the soluble N-terminal and C-terminal domain can collapse in compact structures within 300 to 500 nanoseconds (supp. video 1). The compact structures are a result of dynamic interaction between unpaired polar and charged amino acids (supp. video 1). Ramachandran plots show that compact structures have dominantly β-sheet form with some transient loop structures (Fig 1 A-B). Dynamic structural changes can be always driven by side chains on six His residues, which at physiological pH, can form contacts with either positive or negative groups [38]. Based on the achieved interactions His residues can rapidly change their pKa values and the protonation status [38]. Densely packed polar and charged amino acids at N-terminal and C-terminal domains also form transient contacts with polar and charged lipid heads (methods) (Fig 1 and 2 A-C, [24,33]). Transient protein-lipid interactions can explain why C99-βCTF-APP structures can be affected by the lipid composition [3,24,33-37]. In sum, MD studies can be used to describe different competing interactions that form dynamic N-terminal and C-terminal domains, in correlation with different structural measurements (PDB: 2LP1, [33]).

We analyzed dimerization starting with two free C99-βCTF-APP molecules that were placed 10 to 30 Å apart and facing each other in different orientations (see methods). When placed together C99-βCTF-APP molecules gradually form dimers driven by the diffusion in the bilayer (Fig. 2, supp. video 1). The two C99-βCTF-APP molecules form many transient contacts before dynamic conformers get trapped in a compact dimer (Fig. 2 A-C, supp. video 1). The transient interactions represent transitions through local energy minima before the two structures lock in a stable dimer (supp. video 1). Calculations have been extended to 20 microseconds, which is well beyond the time that takes protein RMSD values to reach a plateau (8 microseconds). About 24.2% of molecular surface area forms an interaction interface in the final complex (Fig. 2 B-C). RMSF values for the individual amino acids [39] show that dimerization is mostly driven by the conformers in the polar parts of the substrate (Fig. 2E). The changes in Lys29-Lys54 distances showed that two C99-βCTF-APP molecules do not have identical structures when they form the dimer, and that dimerization depends on combined contribution from the extended and compact structures (Fig. 2 D). The final dimer structure can be stabilized with as much as 11 H-bonds, and 94 Å^2^ of total interaction surface (Fig. 2 F). Changes in the number of H-bonds as a function of time, show a stepwise increase in the number of H-bonds, as they reflect a sequence of structural changes that drive gradual complex buildup (Fig. 2F).

In conclusion, different structural studies showed that C99-βCTF-APP are highly dynamic molecules that have numerous conformers, which can be difficult to measure, and can be readily affected by the experimental conditions [3,24,33-37]. We show that multiscale MD studies can fill the gaps and trace different structural features of C99-βCTF-APP molecules down to atomic details (Fig. 1 and 2, supp. video 1).

### Multiscale MD studies of saturation of γ-secretase with its C99-βCTF-APP substrate

We analyzed to what extent the interactions observed between two free C99-βCTF-APP molecules (Fig. 2), can be observed when one molecules is bound as a substrate molecule to γ-secretase (Fig. 3 A-C). Specifically, a free C99-βCTF-APP molecule is used to challenge γ-secretase while the enzyme is processing its Aβ substrate (methods). γ-Secretase can be simultaneously exposed to two substrate molecules when the enzyme is gradually saturated with its substrate [12,17,18], i.e. when C99-βCTF-APP substrate start to accumulate next to γ-secretase while the enzyme is still processing its substrate (Fig. 3 A-C, supp. video 2). Some studies indicated that γ-secretase has a separate substrate docking site and the active site [9,30,32], or that γ-secretase can bind multiple substrate molecules in parallel [9,10,12].

**Figure 3.**
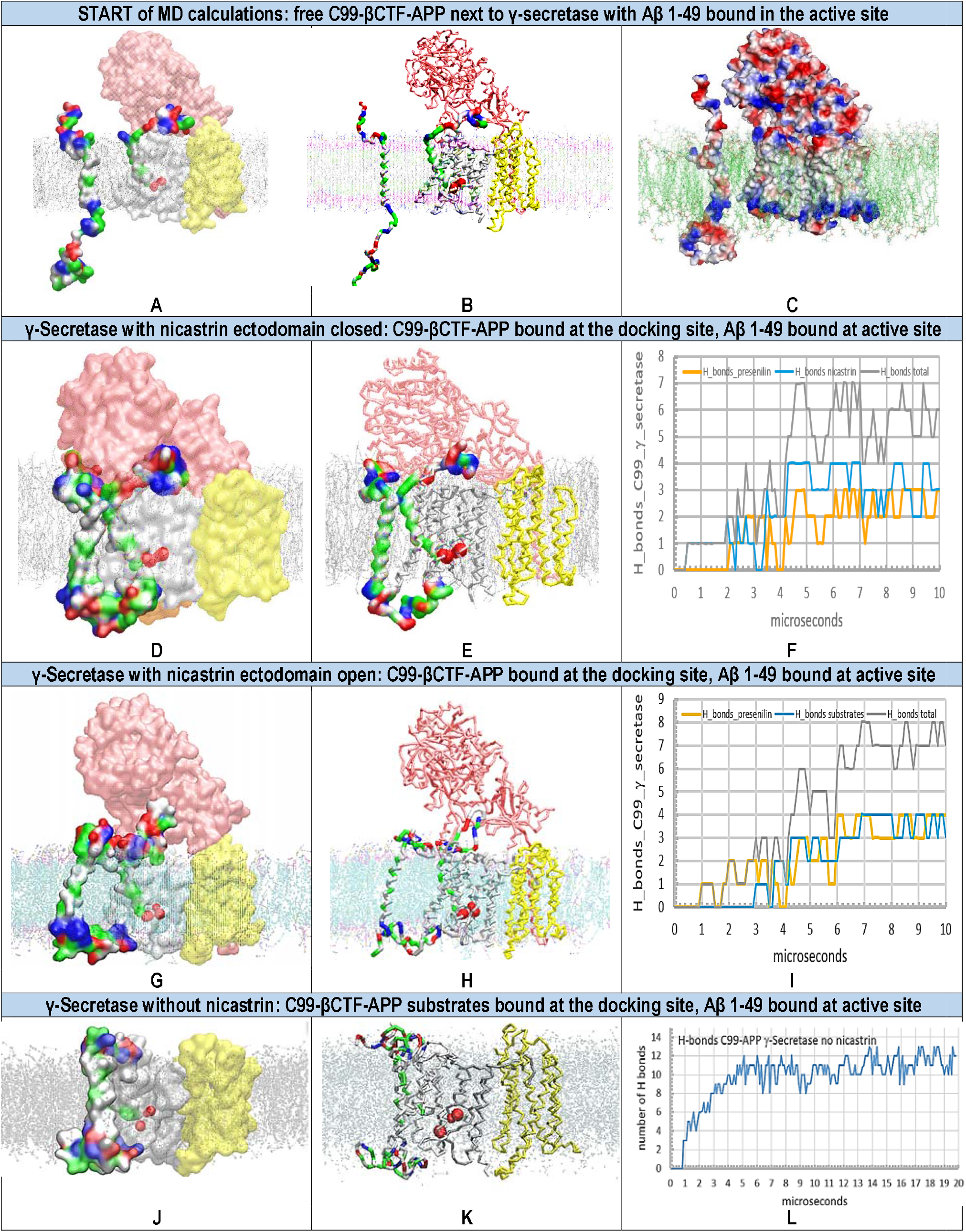
(<u>A</u>-<u>L</u>). Multiscale MD studies of docking of free C99-βCTF-APP substrate to γ-secretase, when its Aβ 1-49 substrate is buried under nicastrin ectodomain. γ-Secretase complex (PDB:6IYC, [32]) is shown as: nicastrin (pink), presenilin 1 (white), Aph1 and Pen2 (yellow). Red blobs depict the active site Asp 257 and Asp 385. The cholesterol-lipid-bilayer is shown as dots (methods). Surfaces of free C99-βCTF-APP (PDB:2LP1, [33]) and bound Aβ 1-49 substrates (PDB:6IYC, [32]) are colored as hydrophobic (white), negative (red), positive (blue), and polar not charged (green). The backbone models are used to show protein conformers, while the partially transparent Connolly surfaces are used to show protein-protein contacts [102]. (**<u>A</u>-<u>B</u>**) MD calculations are designed to represent the increase in the saturation of γ-secretase with its C99-βCTF-APP substrate. Thus, all MD calculations started with free C99-βCTF-APP substrate placed in different rotations 5 to 30 Å away from γ-secretase-(Aβ 1-49) complex. The saturation leads to increased chances that free C99-βCTF-APP substrate can challenge γ-secretase while the enzyme is still processing its Aβ 1-49 substrate [10,17,18]. Multiscale MD studies can describe diffusion in the cholesterol-lipid bilayer, possible docking between the proteins, and the related conformational changes in protein structures (supp. video 2). (**<u>C</u>**) Coulombic coloring of surfaces of γ-secretase and C99-βCTF-APP substrate show that the molecules have charged sites on possible interaction surfaces ([103], supp. Fig. 1). Red is negative, blue is positive, and white is not charged. (**<u>D</u>-<u>F</u>**) The first contact is observed between the N-terminal domain of free C99-βCTF-APP and nicastrin ectodomain as indicated in the past [21-23]. In the closed position, nicastrin ectodomain can affect access of the C-terminal domain of free C99-βCTF-APP substrate to presenilin 1 and Aβ 1-49 substrate in the active site. The figure shows transient contacts between the C-terminal domain and TM2, TM3, TM6a sites in presenilin 1 that can be observed with some conformers. (**<u>G</u>-<u>I</u>**) The mobility of nicastrin ectodomain can be restricted in MD studies in an open position (POSRES @1000 pN) [27]. With the nicastrin head open, the free C99-βCTF-APP substrate can dock with its full length to presenilin 1 in the first contact. The C99-βCTF-APP bound at the docking site can even form contacts with the N-terminal domain of the bound Aβ 1-49 substrate. These results show that nicastrin ectodomain can control docking of the second substrate [21-23]. (**<u>J</u>-<u>L</u>**) When the nicastrin domain is removed from γ-secretase, the free C99-βCTF-APP substrate can dock with its full length to presenilin 1. Its N-terminal can form interactions with the bound Aβ 1-49 substrate. The large interaction surface results in the highest number of H-bonds, while the compact complex structure makes the two substrates indistinguishable. In all cases, the docking can be described quantitatively by following H-bonds as a function of molecular time. The initial lag represents free diffusion and the first contact, while the stepwise changes in the number of H-bonds represent gradual conformational changes in the build-up of the complex.

The parts of C99 structures that support formation of C99 dimers (Fig. 2, supp. video 1), are not visible in the cryo-EM structures of γ-secretase [32] (methods). Thus, these structures are highly mobile, and exist in multiple conformations, even when the substrate is bound and covalently fixed to the γ-secretase [32]. Possible conformers can be predicted by multiscale molecular dynamics studies [27]. We started MD studies with Aβ substrate buried below nicastrin ectodomain, and with its N-terminal domain fully unfolded, just as in the studies with free C99-βCTF-APP substrate (Fig. 3 A-C, supp. video 2). Specifically, we focused first on γ-secretase in complex with Aβ 1-49 substrate, which could represent one of the key steps in pathogenic changes in Aβ production [13,25,40].

Coarse-grained MD studies started with free C99-βCTF-APP substrate placed in cholesterol-lipid-bilayer, facing γ-secretase with Aβ 1-49 bound in the active site (Fig. 3 A-C, methods). We find that nicastrin ectodomain can gradually close over presenilin and the N-terminal of the bound substrate within 2 microseconds (Fig. 3 D-I, supp. Fig. 2 and video 2). The closing was described in the earlier studies [21-23], all of which, have suggested that the closing has several regulatory functions. We find that the closing is driven by interactions between nicastrin, presenilin 1, and presenilin enhancer 2 subunits (described to atomic details in supp. Fig. 2). The closing of the nicastrin ectodomain takes place in parallel with the formation of first contacts between the free C99-βCTF-APP substrate and the nicastrin ectodomain (supp. video 2).

The free C99-βCTF-APP substrate can diffuse and form contacts with γ-secretase driven by complementary electric fields (supp. Fig. 1). It appears, that in its closed position nicastrin ectodomain can affect the free substrate in reaching presenilin 1 (Fig. 3 D-E). The first contacts always form between the N-terminal domain of the free substrate and the nicastrin ectodomain domain, just as suggested in the earlier studies [22]. Subsequently the C-terminal starts to form transient contacts with TM2 and TM3 on presenilin 1 (supp. video 2).

We further explored possible functions of nicastrin ectodomain by repeating the CG-MD calculations with the nicastrin head fixed in open position (Fig. 3 G-I, posrestrain 1000pN [27])). The restrained nicastrin does not close in calculations that represent as much as 20 microseconds of molecular time (Fig. 3 G-H). When the nicastrin ectodomain is open the free substrate can immediately form contacts with Aβ substrate and presenilin (Fig. 3 I). We observe in the first contact as many as 6 hydrophobic interactions, and up to 5 polar interactions (Fig. 3, G-I). Such interactions can readily affect dynamic structures that control catalytic functions around TM2, TM3, TM6a, and Aβ substrate [25,32]. The regulatory function of nicastrin ectodomain is further demonstrated by repeating the multiscale MD studies with γ-secretase without nicastrin (Figs. 3 J-L). In the absence of interference by nicastrin, the free C99-βCTF-APP substrate can make tight interactions with the bound substrate and presenilin 1 with its full length (Figs. 3 J-L).

In conclusion, we have presented a novel two-substrate mechanism which shows that the second substrate can dock to γ-secretase while the enzyme is still processing its Aβ substrate (Fig. 3, supp. video 2). The presented two-substrate mechanism has multiple significant consequences. The second substrate binds to the most dynamic γ-secretase structures that control Aβ production [8,20,25,32]. Thus, presented two-substrate mechanism can explain why saturation of γ-secretase with its substrate leads to pathogenic changes in catalytic activity [10,11,13,14]. The presented two substrate mechanism is an extension of the earlier studies which showed that γ-secretase has a separate substrate docking site and the active site [9,16,26,30,32]. The presented two substrate mechanism is an extension of the earlier studies which have suggested that nicastrin ectodomain can control substrate docking [21-23].

### Multiscale MD studies of nicastrin function in γ-secretase complex with the exposed N-terminal end of the bound Aβ substrate

When an Aβ substrate is bound in the active site of γ-secretase, its highly mobile N-terminal domain can be hidden to a different degree under nicastrin ectodomain [32] (see methods). In one extreme, the N-terminal domain can be fully exposed at the external surface of nicastrin ectodomain (see methods, Fig. 4A). We have prepared γ-secretase in a complex with Aβ 1-49 substrate that had the N-terminal domain exposed on the nicastrin external surface (Fig. 4A). The complex was challenged with free C99-βCTF-APP substrate. The aim was to analyze to what extent the nicastrin ectodomain can prevent potentially toxic aggregation between N-terminal domains of the two substrates (i.e. to compare Figs. 3 and 4).

**Figure 4.**
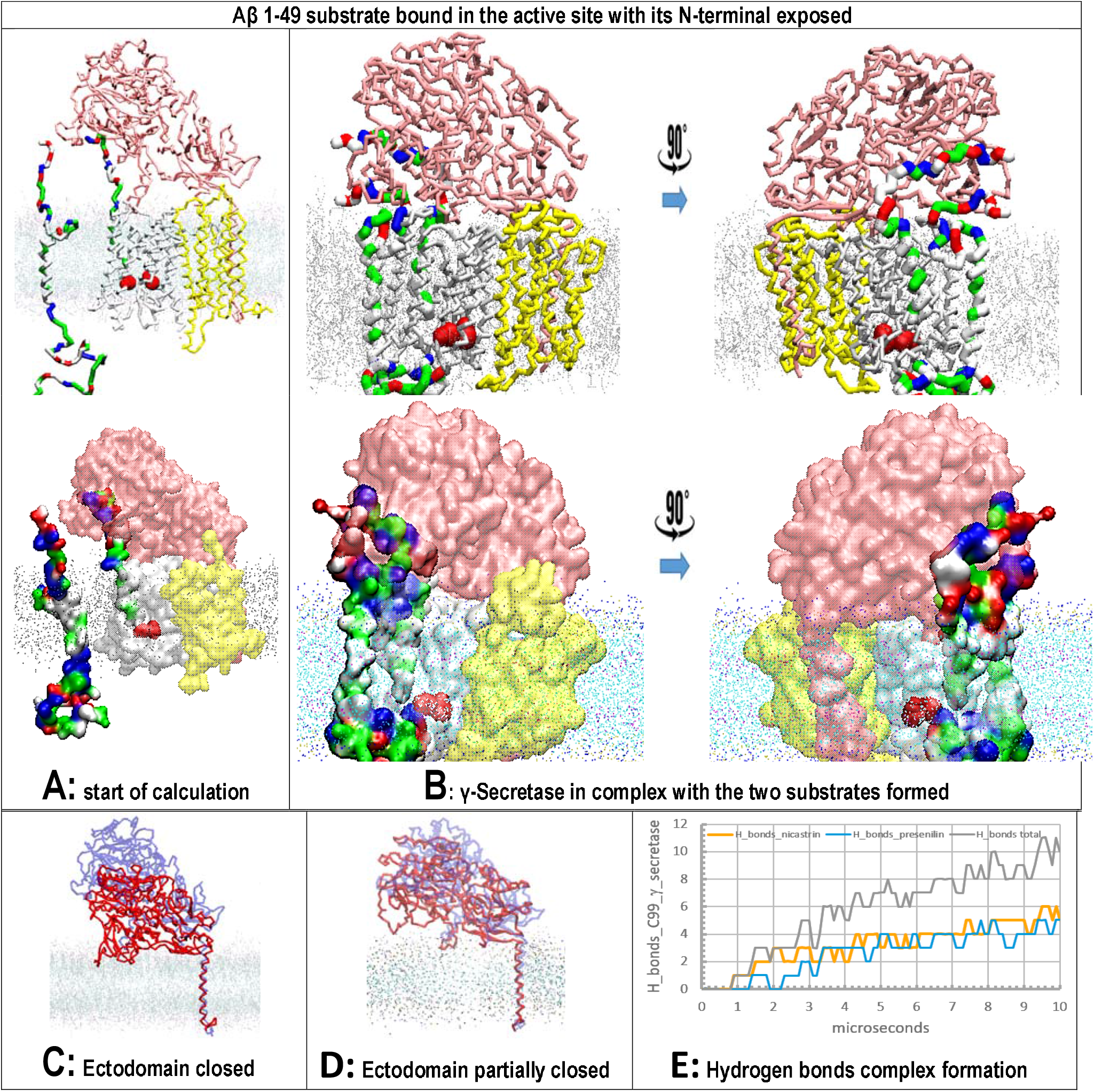
(<u>A</u>-<u>E</u>). Multiscale MD studies of docking of free C99-βCTF-APP substrate to γ-secretase, when its Aβ 1-49 substrate is exposed on the nicastrin external surface. γ-Secretase complex (PDB:6IYC, [32]) is shown as: nicastrin (pink), presenilin 1 (white), Aph1 and Pen2 (yellow). Red blobs depict the active site Asp 257 and Asp 385. The cholesterol-lipid-bilayer is shown as dots (methods). Surfaces of free C99-βCTF-APP (PDB:2LP1,[33]) and bound Aβ 1-49 substrates (PDB:6IYC, [32]) are colored as hydrophobic (white), negative (red), positive (blue), and polar not charged (green). The backbone models are used to show protein conformers, while the partially transparent Connolly surfaces are used to show protein-protein contacts [102]. (**<u>A</u>**) at the start of calculations, the free C99-βCTF-APP substrate is positioned 5 to 30 Å away from γ-secretase in complex with its Aβ 1-49 substrate. Just as in figures 3 (A-B), the nicastrin ectodomain is open at the start of calculation [21-23], but this time, the N-terminal of the bound Aβ 1-49 substrate is fully exposed on the nicastrin surface. (**<u>B</u>**) The free C99-βCTF-APP can diffuse through the membrane and dock to γ-secretase-(Aβ 1-49) complex. N-terminal parts of two substrates compete in interactions with the nicastrin ectodomain. TM parts of the two substrates form binding interactions at the start of the active site tunnel (sites between TM2 and TM3). C-terminal part of the free C99-βCTF-APP forms transient interactions with presenilin at its most dynamic cytosolic sites, the end to the active site tunnel [25,32]. (**<u>C</u>**<u>-</u>**<u>D</u>**) nicastrin ectodomain at the start (blue) and the end (red) of MD calculation. The models on the right show that exposed N-terminal of Aβ 1-49 substrate can affect the closing of the nicastrin ectodomain. The closed ectodomain can take many conformers that can be difficult to capture by cryo-EM studies [32]. (**<u>E</u>**) H-bonds as a function of the calculated time depict a gradual buildup of the complex. The lag time represents the initial diffusion and the first contact, while the stepwise changes in the number of H-bonds represent gradual conformational changes in the build-up of the complex.

Multiscale MD studies show that the N-terminal domain of the substrate is highly flexible and always in contact with the nicastrin surface (Fig. 4A). Different conformers always lead to some binding interactions because both proteins share numerous polar and charged groups on their surface (Fig. 1C, and supp. Fig. 1). The N-terminal domain of the bound substrate cannot prevent the closing of the nicastrin ectodomain but it can affect related conformational changes (Fig. 4 C-D).

The docking of the free C99-βCTF-APP substrate is not significantly affected by the position of the N-terminal domain of the bound substrate (compare Fig. 4 and Fig. 3 E-F). Most notably, the nicastrin ectodomain can bind the N-terminal domains of both substrates, and thus, compete with potentially toxic aggregation between those molecules (compare Fig. 4B with Fig. 3 I-F). We are proposing that the nicastrin ectodomain can prevent toxic aggregation between the N-terminal domains of the two substrates by different mechanisms in different conformations (compare Fig. 3-4). Closed nicastrin ectodomain can take multiple conformations in its function (Fig. 4 C-D), what can explain why cryo-EM studies could not capture nicastrin structure in closed conformations [32].

### Multiscale molecular dynamics studies of γ-secretase with the two substrates of different lengths

γ-Secretase can use substrates of different lengths, that come from α-secretase and β-secretase reaction paths: C83-αCTF-APP and C99-βCTF-APP [32]. The substrate length can affect the pathogenic changes in Aβ metabolism [41]. Multiscale MD studies were used to analyze interactions between γ-secretase and the substrate of different lengths (Fig. 5). Transmembrane section of the free substrate (starting Val12-His13-His13-Gln15; ending with Lys53-Lys54-Lys55-Gln56-Tyr57) was used to challenge γ-Secretase in complex with Aβ catalytic intermediates with the shortest N-terminal (starting at Val17).

**Figure 5.**
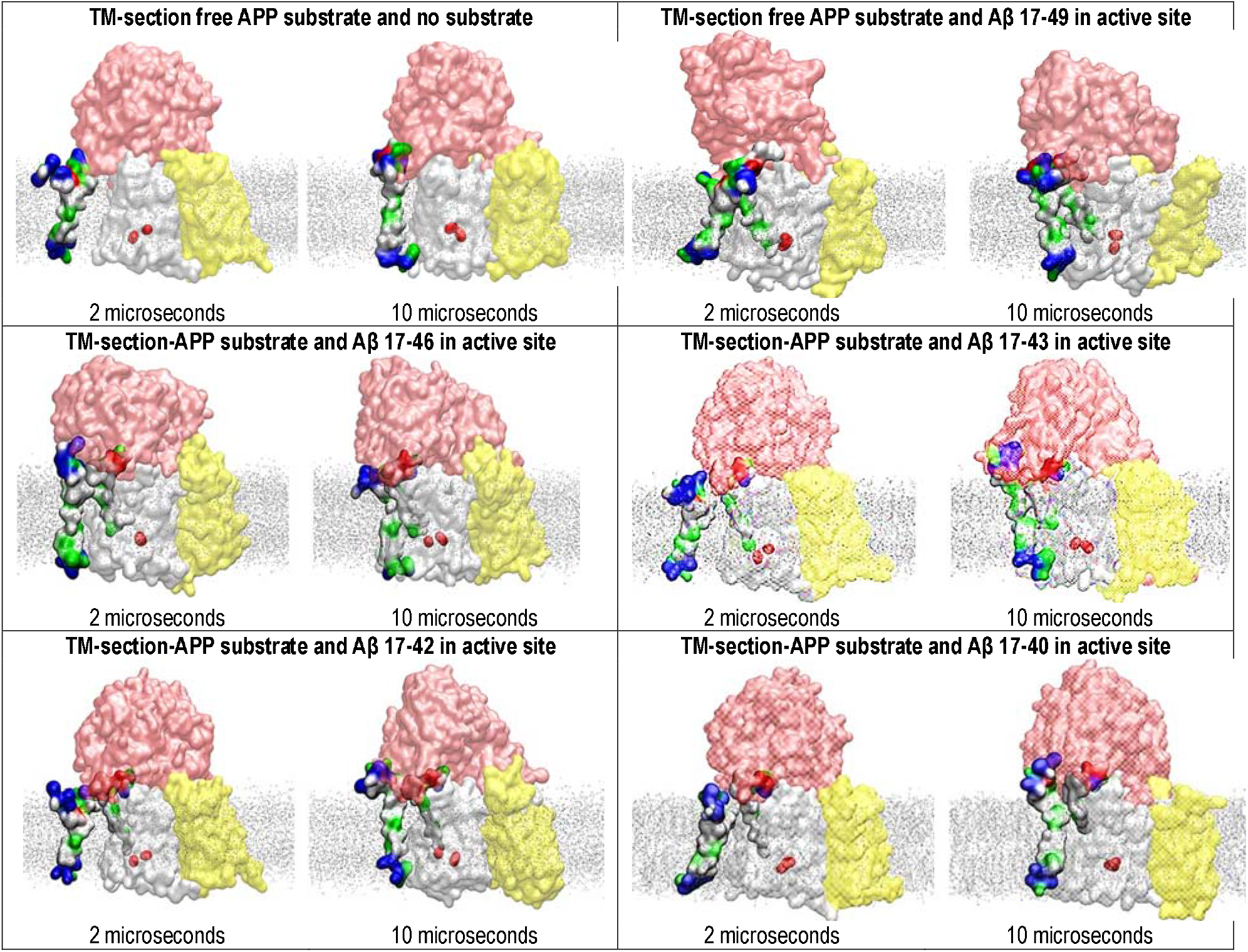
Multiscale MD studies of docking of the shortest substrates forms to γ-secretase, when shortest substrates are bound in the active site (PDB:6IYC, [32]). γ-Secretase complex is depicted as partially transparent Connolly surfaces, showing: nicastrin (pink), presenilin 1 (white), Aph1 (yellow), and Pen2 (not visible) [102]. Red blobs depict the active site Asp 257 and Asp 385. The surfaces of the two substrates are shown as hydrophobic (white), negative (red), positive (blue), and polar not charged (green). The cholesterol-lipid-bilayer is shown as silver dots. The calculations used the shortest substrate forms to depict the minimal effect that substrate binding at the docking site can have on the substrates bound at the active site. The free substrate shows the NMR structure, starting V12-H13-H13-Q15, ending with 53K-54K-55K-Q56-57Y (PDB:2LP1, [33]). Th substrate bound in the active site tunnel had different lengths Aβ 17-x, to represent different catalytic intermediates that can form from 83-α-CTF-APP substrate [11,13]. The short N-terminal of the bound substrate is hidden under nicastrin ectodomain [32]. We compared the structures at the first contact between the free substrate and γ-secretase (at 2 microseconds), with the structures of the fully formed complex (at 10 microseconds). The figures show that even the shortest substrates can form contacts that can affect dynamic changes in presenilin structure that drive processive catalysis [13].

We found that in all cases two substrates molecules can bind simultaneously to γ-secretase (Fig. 5), however, the shorter substrates can make a smaller number of binding interactions (compare H-bonds Figs. 3, 4, and 5). Thus docking of the shorter free substrate is less likely to affect γ-secretase activity. These results are consistent with the different studies which showed that shortest substrates are less likely to support pathogenic changes in Aβ products [41]. Comparative analysis of docking mechanisms with different substrates demonstrates the power of multiscale MD studies in describing the presented interactions (Fig. 5).

### Multiscale molecular dynamics studies of docking of free C99-βCTF-APP substrate to γ-Secretase complex with no substrate-bound (Fig. 6)

γ-Secretase complex with no substrate-bound was challenged with free C99-βCTF-APP substrate in different orientations (Fig. 6). In the absence of the bond substrate nicastrin ectodomain can completely close over presenilin 1 (Fig. 6, supp. Fig. 2). The free C99-βCTF-APP substrate can diffuse in the membrane and bind to the nicastrin ectodomain because both proteins share numerous polar groups (Fig. 6, supp. Fig. 1). The closing of the nicastrin ectodomain can also interfere with the contacts between the free C99-βCTF-APP substrate and presenilin 1 (Fig. 6 B-D). The free substrate can dock on the nicastrin head but could not reach the presenilin surface (Fig. 6C). These results further support the idea that closing of the nicastrin ectodomain can be a significant mechanism in the regulation of substrate docking to γ-secretase [21-23]. The closing mechanism can be a possible target for future drug-design efforts, specifically for the development of competitive inhibitors of γ-secretase that can decrease the enzyme saturation with its substrate like the protective mutation [41].

**Figure 6.**
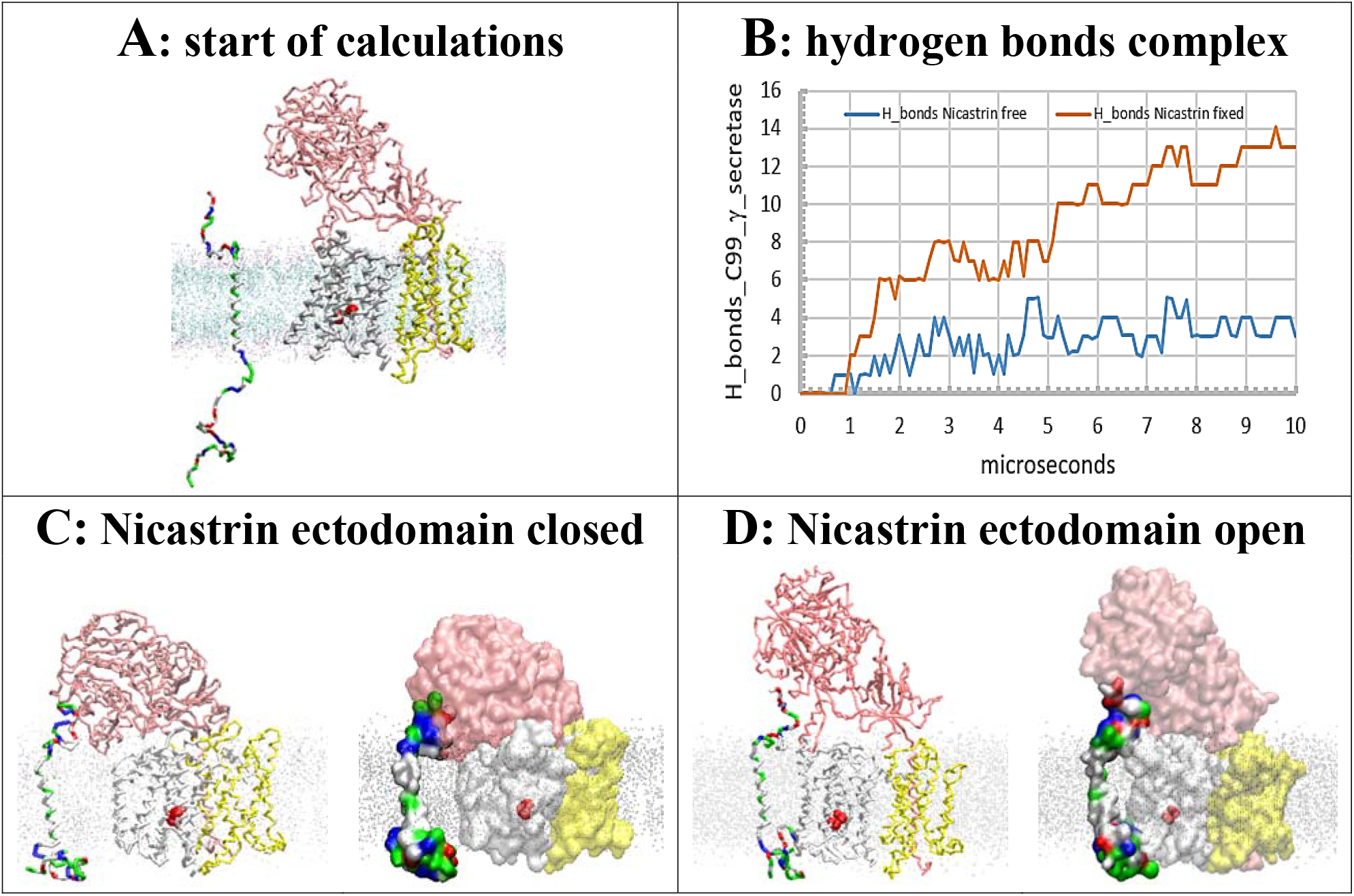
(<u>A</u>-<u>D</u>). Multiscale molecular dynamics studies of nicastrin function in the regulation of docking of free C99-βCTF-APP substrate to γ-secretase with no substrate bound in the active site. γ-Secretase complex is shown as: nicastrin (pink), presenilin 1 (white), Aph1, and Pen2 (yellow). Red blobs depict the active site Asp 275 and Asp 385. The silver lines represent cholesterol-lipid-bilayer. Free C99-βCTF-APP substrate is colored as hydrophobic (white), negative (red), positive (blue), and polar not charged (green). The backbone models are used to show protein conformers, while the partiall transparent Connolly surfaces are used to show protein-protein contacts [102]. (**<u>A</u>**) Residue based coarse-grained MD calculations started with the fully extended C99-β-CTF-APP structure that was positioned 20 Å away from γ-secretase with the nicastrin ectodomain open. (**<u>B</u>**) The buildup of protein-protein interactions is described quantitatively by counting H bonds that form in calculated molecular time (10 microseconds). The initial lag represents the diffusion time and the conformational changes that take place after the first contact. The subsequent steps in the graph represent conformational changes in the complementary surfaces that drive the buildup of binding interactions. (**<u>C</u>**) In the closed position nicastrin ectodomain can block access of the free C99-βCTF-APP substrate to TM2 and TM3 sites in presenilin 1. A maximum of 3 to 4 H bonds is observed when the free nicastrin head falls and blocks the substrate from reaching presenilin, or making a full set of contacts with presenilin 1. (**<u>D</u>**) The mobility of nicastrin ectodomain can be restricted in MD studies in an open position (POSRES @1000 pN) [27]. With the nicastrin head open the free C99-βCTF-APP substrate can dock with its full length to presenilin 1 and nicastrin. The docking sites on presenilin overlap with dynamic protein structures that can be affected by drugs and FAD mutations [8,32], and can affect Aβ production and catalytic efficiency [32]. As many as 13 hydrogen bonds can be observed when the substrate is extended over the full length of the γ-secretase complex.

In conclusion, we have presented our two-substrates mechanism in four very different situations (Figs. 3-6). First, with one substrate fully buried under nicastrin ectodomain (Fig. 3, supp. video 2). Second, with one substrate exposed on the surface of the nicastrin ectodomain (Fig. 4). Third, with the substrates of different lengths (Fig. 5, supp. Fig. 3). Forth, with no substrates bound at the active site tunnel (Fig. 6). These are just some of the selected interactions, out of many intermediate situations, which we could not show due to respect for the space limitations. Different interactions show differences in the rate of contact buildup, in the contact sites, and the final number of H-bonds formed (Figs. 3-6). Nevertheless, all four presented situations support several major conclusions. In all cases, the N-terminal of the free substrate makes contacts with the nicastrin ectodomain (Figs. 3-6) [22]. Those contacts can control substrate docking to γ-secretase to a different extent (Figs. 3-6). In all four cases, the initial interaction by the N-terminal domain controls the contact between the C-terminal domain and the cytosolic end of presenilin 1 (Figs. 3-6). Possible contacts between the C-terminal domain and cytosolic end of presenilin 1 are very significant. Those sites can control processive catalysis, and those sites can bind drugs and get affected by FAD mutations [8,20,25,32].

### AA-MD studies of docking of C-terminal domain of C99-βCTF-APP to cytosolic section of presenilin subunit

So far we learned that docking of the N-terminal domain of free C99-βCTF-APP substrate to nicastrin leads to gradual docking of its C-terminal to presenilin (Figs. 3-5, supp. video 2) [21-23]. The C-terminal domain is docking to the most dynamic parts in presenilin structure that control processive catalysis [25,32]. Thus we have analyzed the docking interactions to atomic details using conversion from CG-MD to AA-MD studies (Fig. 7, and supp. video 3). AA-MD studies started with CG-MD structures that represent the first contacts between the C-terminal domain of and the cytosolic end of presenilin 1 (CG-MD to AA-MD conversion see methods). Different calculations show some differences in docking sites and the related structural changes in active site tunnel on presenilin subunit [25]. The differences are due to different contact surface size, or due to different docking orientations, or due to protonation of the two active site Asp, or due to lipid composition of the membrane (supp. Fig 3, and manuscript in preparation). However, we also found some common features in all of our docking calculations (Fig. 7).

**Figure 7.**
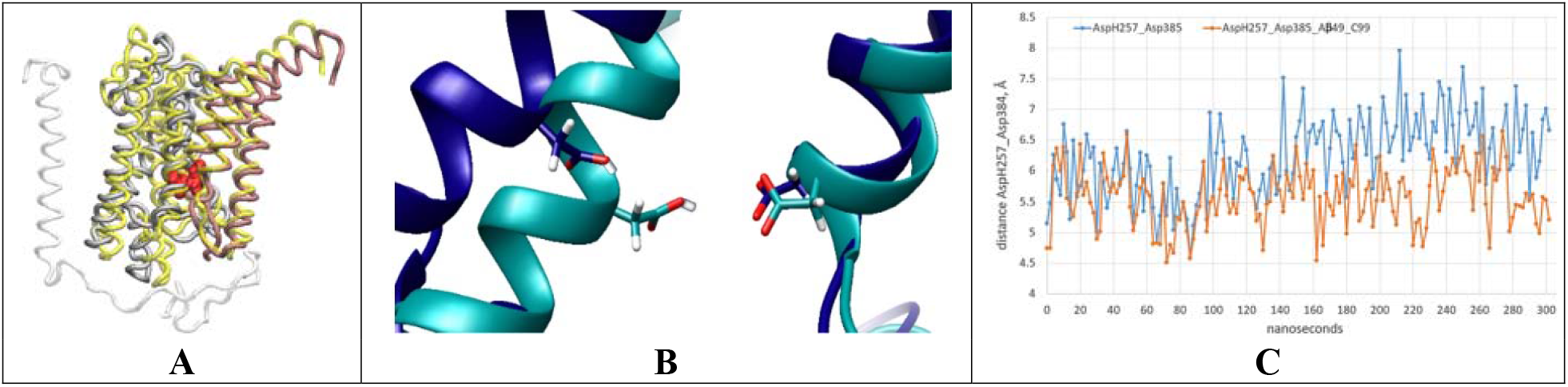
(<u>A</u>-<u>C</u>). AA-MD studies of docking interactions between C-terminal section of C99-βCTF-APP substrate and cytosolic section of γ-secretase-(Aβ 1-49) complex [32]. (**<u>A</u>**) for clarity the figure is focused only on parts that participate in the interaction (Fig. 3D, supp. video 3). Namely presenilin structure (gray) with C99-βCTF-APP fully docked (white) is superimposed on presenilin structure before the docking (N-terminal domain yellow, C-terminal domain pink) [32]. The superimposed structures show that C99-βCTF-APP bound at the docking site can spread apart cytosolic ends of TM2, TM3, TM6 and TM6a by acting on the loops between the TM regions. The spread-out can affect key parts in the processive catalysis [25] including the active site Asp257 and Asp385 (shown in red). The docking affects presenilin structure at the sites that are most frequently affected by FAD mutations [25,32] and drugs [8]. (**<u>B</u>**) docking of free C99-βCTF-APP to cytosolic end of presenilin subunit leads to increase in angle and distance between the active site Asp257 (protonated) and Asp385 (unprotonated). The changes can affect optimal catalytic structures of γ-secretase [25] (**<u>C</u>**) AA-MD analysis of changes in distance between γ-carbon atoms on AspH257 and Asp385 caused by C99-βCTF-APP docking as a function of the calculated time. We compared the changes in AspH257 and Asp385 distances caused by the docking (blue) with the changes in the absence of C99-βCTF-APP (red).

In all calculations the docking of the substrate C-terminal end will gradually spread apart the cytosolic ends of TM2, TM3 and TM6 on presenilin subunit by acting on the connecting structural loops and TM6a (Fig. 7A, supp. video 3). The spreading will result in the opening of the active site tunnel and increase in distance and angle between the active site Asp257 and Asp385 (Fig. 7 B-C). The docking affects presenilin structure at the sites that are most frequently affected by FAD mutations [25,32,42]. The docking also affects known drug-binding sites [8,20]. The docking can increase the average distance between active site Asp257 and Asp385, just like the FAD mutations (supp. Fig. 3), or binding of drugs [8], or a switch from POPC-bilayer to cholesterol-lipid-mix-bilayer (supp. Fig. 3).

In conclusion, we propose that docking of C-terminal of C99-βCTF-APP substrate to the cytosolic end of presenilin subunit could explain, at the molecular structure level, why saturation with the substrate leads to changes in the of γ-secretase activity [6,9,10,13,14,40]. Namely, the shifts from Aβ(x-49) to Aβ(x-48) production [10,13,40], increase in the Aβ(x-42)/Aβ (x-40) ratio [13,40], and changes in the enzyme’s response to different-drugs [6,9].

### Substrate channeling between BACE1 and γ-secretase

Presented results are suggesting that the nicastrin ectodomain can be the key mechanism that can control binding of the second substrate to catalytically active γ-secretase (Figs. 3-6). In that aspect the presented two-substrate mechanism is an extension of the earlier studies [21-23]. In cells opening-and-closing of nicastrin ectodomain could be regulated by interaction between γ-secretase and β-secretase (BACE1) [43]. Such interactions would indicate that substrate channeling can regulate substrate docking to γ-secretase, and thus, disease pathogenesis [44]. Regulation of enzyme activity by substrate channeling is frequently observed in metabolic studies [45]. The proteins in cells are present in exceptionally high concentrations that favor supramolecular complexes [45]. We used multiscale MD studies to analyze docking of β-secretase to the nicastrin ectodomain (Fig. 8, [43]).

**Figure 8.**
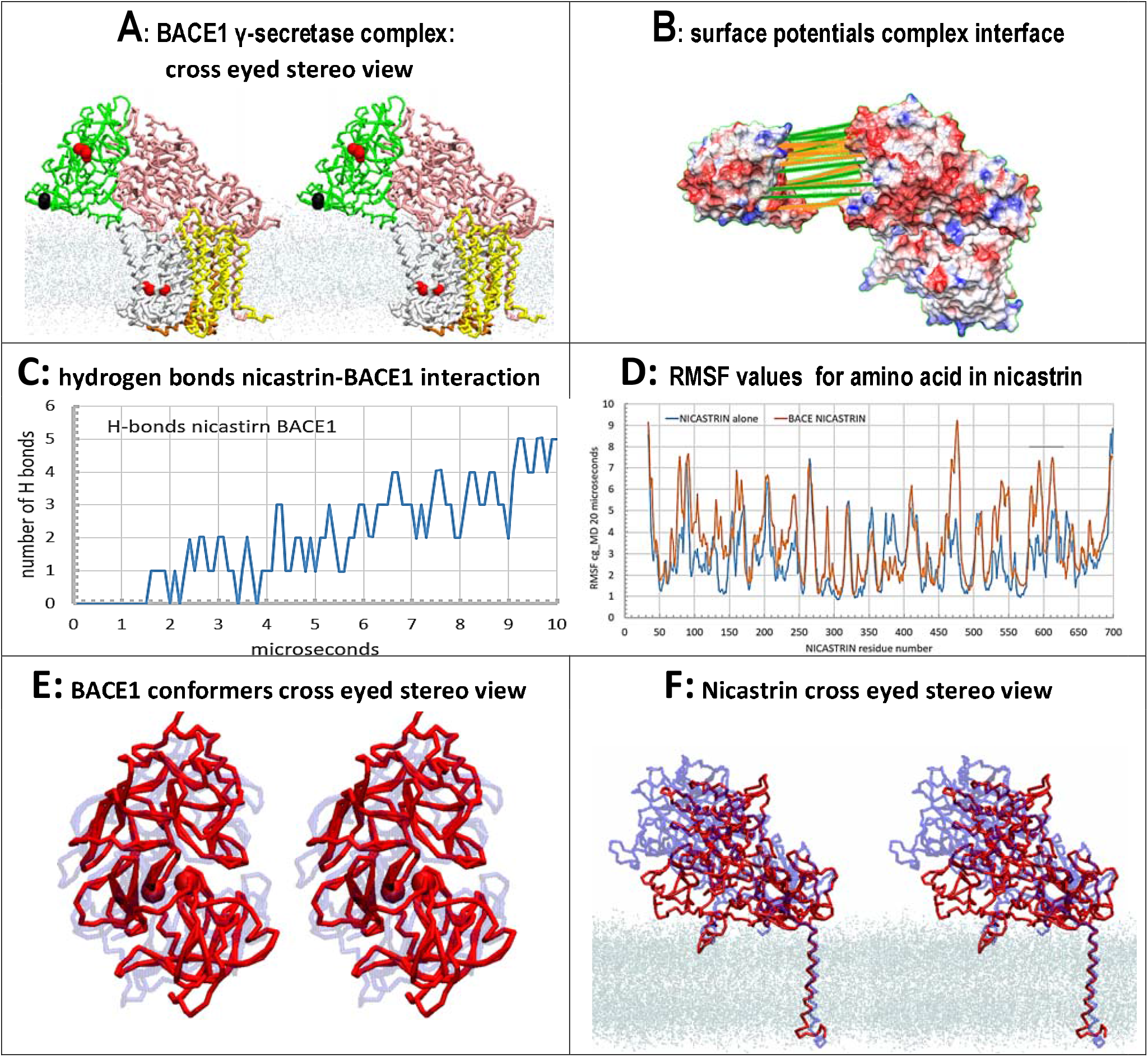
(<u>A</u>-<u>F</u>). Multiscale MD studies of complex between BACE1 (PDB:4FGX, [46]) and γ-secretase (PDB:6IYC, [32]). (**<u>A</u>**) Cross-eyed stereo presentation is used to maximally highlight conformational changes induced by complex formation. BACE1 (green) is shown with highlighted actives site aspartates (red) and C-terminal (black). γ-Secretase shows nicastrin (pink), presenilin (silver), Aph1 (yellow), and active site Asp 257 and Asp 385 as red vdw spheres. Cholesterol-lipid-bilayer is shown as dots. (**<u>B</u>**) Proteins are shown as Connolly surfaces with the interacting surfaces spread apart to show the matching surface shapes and the electric potentials. The surfaces are colored according to charges of surface amino acids, positive (blue), negative (red), neutral (white) [103], with highlighted H-bonds sites (green) and vdw contacts (gold). (**<u>C</u>**) The complex build-up in MD calculations was monitored by counting H-bonds formed between the proteins. The initial lag represents diffusion and the first contact, while the stepwise increase in the number of hydrogen bonds represents conformational changes that drive complex formation. (**<u>D</u>**) comparative analysis of RMSF values [39] can be used to analyze to what extent different parts of nicastrin structure can get affected by formation of the complex. (**<u>E</u>**<u>-</u>**<u>F</u>**) Repeated docking showed that the biggest interaction surface and the largest number of hydrogen bonds can be seen when the active site cleft of BACE1 is open (E) and the nicastrin ectodomain is adapted for interaction (F). Cross-eyed stereo presentation is used to maximally highlight conformational changes induced by complex formation. The conformational changes are illustrated by overlapping BACE1 and nicastrin ectodomain structure before the complex formation (blue) and after the complex formation (red). Cholesterol-lipid-bilayer is shown as dots.

When β-secretase was positioned facing the nicastrin ectodomain in the plane of TM2 and TM3 the two proteins form transient interactions with their polar surfaces (Fig. 8). The specificity of the presented docking was tested by doing a series of docking studies. β-secretase was placed 5 to 15 Å away from nicastrin, facing different sites, at different angles. Different setup for the initial docking interface can affect the rate of complex formation, due to a competition between the docking interactions and the closing of the nicastrin ectodomain (Fig. 8F). Furthermore docking studies intentionally used β-secretase structure that does not have transmembrane domains (PDB:4FGX, [46]). This structure can readily diffuse in the surrounding media, unless it forms binding interactions. β-secretase in presented complex is docked to nicastrin with its C-terminal facing membrane surface (Fig. 8A, black spheres). This is expected orientation for start of transmembrane domain for β-secretase structure [46].

The biggest interaction surface was observed when the nicastrin ectodomain was open, and facing β-secretase in plane that is parallel to TM2 and TM3 plane on presenilin 1 (Fig. 8A and 8F). These interactions can induce structural changes in both β-secretase and γ-secretase, which can further facilitate buildup of interaction sites (Fig. 8B). β-secretase can embrace nicastrin by opening its active site loops, residues loops 94-97 and loop 158 to 162, and loops 358-365 and loops 317 to 322 (Fig. 8E) [46]. Nicastrin ectodomain will mold to β-secretase structure with its highly dynamic β -sheet structures (residues 576-623). Interactions domain is 576 to 623 with sugar on Asn580. The contact sites are Asp264, Thr268, and Arg270, and on Gln585, Ser594 and Asn596 (Fig. 8B). β-secretase interaction contacts are on Asn159, Gln319, Glu358, Ser363, Asp365 [46] (Fig. 8B).

Interaction with BACE1 is pushing nicastrin ectodomain in opposite direction from the conformational changes that take place when the ectodomain is guarding substrate access to the docking site (compare Fig. 8A with Fig. 6C). The complex between BACE1 and γ-secretase is stretching the flexible and long loop between TM2 and TM1, and thus, opening the space between TM2 and TM3 (Fig. 8A). Closing and opening of the substrate binding sites is known to regulate transient protein-protein interactions in the case of substrate channeling [45].

## Discussions and Conclusions

We will demonstrate the significance of the presented two-substrate-mechanism by showing that this mechanism can address a wide range of observations from different studies of Alzheimer’s disease. Numerous studies have suggested that γ-secretase has separate substrate docking site and active site [7,9,10,16,30,32,47]. Here we go a step further. We show that γ-secretase can bind in parallel two different substrate molecules, one at the docking site and one at the active site (Figs. 3-5). Two substrates can bind to γ-secretase when the enzyme is gradually saturated with its substrate (Fig. 9). In such conditions, free C99-βCTF-APP can start to accumulate next to γ-secretase while the enzyme is still processing its first substrate (Fig. 3 A-C, supp. video 2). The physiological significance of the presented two-substrate-mechanism will be summarized around five closely related observations from different studies of changes in γ-secretase activity in Alzheimer’s disease. Development of early diagnostic methods and effective drug-design strategies depends on our ability to connect observations from different enzyme-based, cell-based, animal-based, and clinical studies of Alzheimer’s disease [3,5,7,9,10,29].

**Figure 9.**
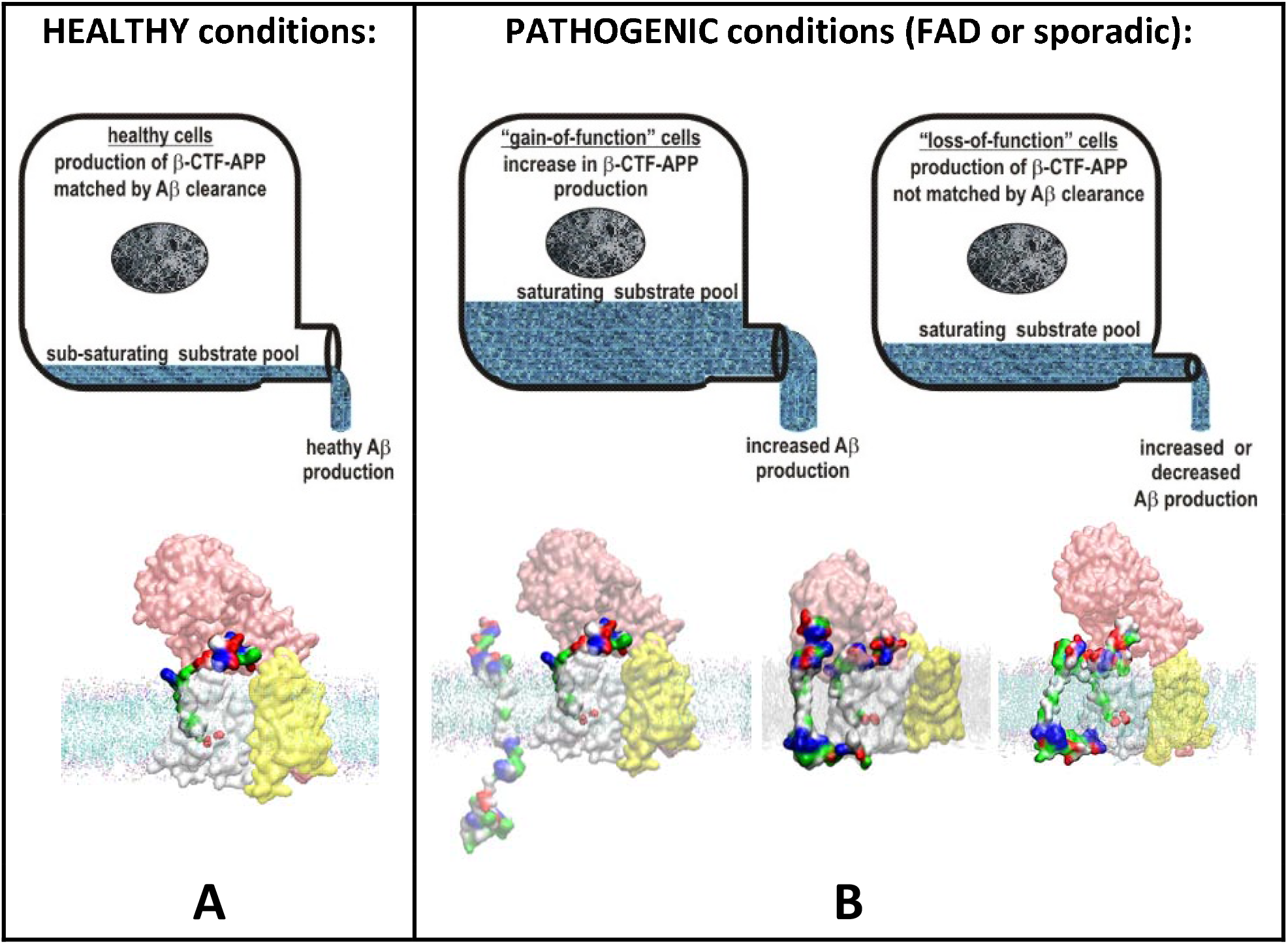
(<u>A</u>-<u>B</u>). Summary figure: decrease in catalytic capacity can trigger pathogenic saturation of γ-secretase in sporadic and familial cases of Alzheimer’s disease. The figure is prepared to summarize the physiological significance of the presented structural analysis and the related studies from the past. γ-Secretase can be illustrated as a drain pipe for cellular amyloid metabolism. Different levels of amyloid metabolism are illustrated as different levels of drain load. (**<u>A</u>**) in healthy cells, γ-secretase can completely process its different C99-βCTF-APP, C83-αCTF-APP, and Aβ substrates to soluble fragments with no interference [13,25,40] The catalytic capacity of γ-secretase can match cellular levels of amyloid metabolism. (**<u>B</u>**) in pathogenic conditions, the catalytic capacity of γ-secretase can NOT match amyloid metabolism. The mismatch can be due to increase in amyloid metabolism (left), or due to decrease in the maximal activity of γ-secretase (right), or due to combination of both effects. Any increase in saturation of γ-secretase with its substrate leads to increase in chances that the second substrate can challenge th docking site while the enzyme is still processing its first substrate [25,32]. Binding of the second substrat can trigger a sequence of potentially toxic events. C-terminal domain of the second substrate can affect dynamic presenilin structures that regulate processive catalysis. The same sites bind different drugs, and get affected by disease causing mutations [8,20,32]. Thus, binding of the second substrate can lead to toxic inhibition of γ-secretase activity and toxic increase in production of the longer, more hydrophobic, Aβ peptides [10,13,14]. Such hydrophobic shifts in Aβ production can support pathogenic protein aggregation and inhibition γ-secretase activity, which can lead to cell death. Toxic aggregation between the N-terminal domains of the two substrates is controlled by nicastrin ectodomain. Such aggregation is more likely to happen with C99-βCTF-APP than with C83-αCTF-APP substrate, which can explain, why β-secretase path is more pathogenic than α-secretase path. Alzheimer’s disease can be described at the molecular structure level as γ-secretase choking on an excess of its sticky substrate.

### 1. The two-substrate mechanism and pathogenic changes in the type of Aβ products

The two most frequently analyzed pathogenic events are an increase in Aβ (x-42)/Aβ (x-40) ratio [13,40] and an increase in the production of the longer more hydrophobic Aβ products [40]. The presented two-substrate mechanism can explain the observed changes in Aβ production (Fig. 7).

A wide range of different studies showed that gradual saturation of γ-secretase with its C99-βCTF-APP substrate leads to an increase in the Aβ (x-42)/ Aβ (x-40) ratio and accumulation of the longer, more hydrophobic, Aβ products [10,13,14]. From basic principles in enzymology, we know that gradual saturation of γ-secretase with its C99-βCTF-APP substrate leads to a gradual increase in the chances that γ-secretase can bind two substrate molecules in parallel [17-19]. Briefly, the catalytic cycle of γ-secretase consists of three steps: substrate recognition-and-binding, catalysis, and product release [17,18]. Substrate recognition-and-binding are limiting step when the enzyme is subsaturated with its substrate [17,18]. The catalysis and the product release are becoming limiting steps when the enzyme is increasingly more saturated with its substrate [17,18]. Thus, the gradual saturation leads to increased chances that the second substrate can challenge the docking site, while the enzyme is still processing its first substrate (Fig. 9).

The second substrate, different drugs, and FAD mutations, target the same dynamic sites in γ-secretase structures (Figs. 3-5 and 7, supp. video 3, [8,20]). These sites control processive catalysis [25,32]. Subtle conformational changes in these sites can induce shift from Aβ 49-46-43-40 path to Aβ 48-45-42 path (Fig. 7, supp. video 3), and thus, increase in Aβ (x-42)/Aβ (x-40) ratio [14,20,25,32,48]. The substrate bound at the docking site can produce partial inhibition or even entrapment of the longer Aβ catalytic intermediates (Figs. 3-5 and 7, supp. video 3). The inhibition and the entrapment can further facilitate saturation with the substrate (Fig. 9, [18]). Thus, the entire process can facilitate the pathogenic events in a positive feedback loop (Fig. 9, [18]).

### 2. The two-substrate mechanism can explain toxic changes in Aβ production in all sporadic and FAD cases of the disease

Both, increase and decrease, in Aβ production can be observed in pathogenic events in Alzheimer’s disease [3,7,49-51]. Thus, the development of effective diagnostic and therapeutic approaches depends on our ability to understand factors that can balance optimal γ-secretase activity and Aβ metabolism (Fig. 9) [3,15]. Any misbalance between γ-secretase activity and Aβ metabolism can lead to accumulation of the substrates, and thus, chances for pathogenic saturation with the substrates (Fig. 9) [3,7,49-51].

All FAD mutations lead to increase in saturation of γ-secretase with its substrate [7,14,52], either by decrease in catalytic rates of γ-secretase [10,11,13,53-55], or by increase in C99-βCTF-APP level [56-58]. In both cases, the end result is an accumulation of different substrates and possible pathogenic interactions (Fig. 9). The earliest age-of-onset can be observed with mutants that have the biggest chance to reach saturation at the lowest substrate loads (Fig. 9) [7]. The protective islandic A673T mutation in APP substrate is the only mutation that leads to a decrease in γ-secretase saturation with its C99-βCTF-APP substrate [41].

Pathogenic processes in sporadic disease can be caused by any age induced decrease in catalytic turnover rates for γ-secretase [15,29,50,59-65], or by any age induced increase in C99-βCTF-APP metabolism [58,66-69], or both processes[7,14,49,50,70,71]. Any of those changes can be driven by age-induced changes in cell physiology [15], or by changes in relative gene expression levels [63,72]. Thus a large number of physiological processes can support development of Alzheimer’s disease [3,49,58,73]. However, all pathogenic processes share the same set of events: decrease in the γ-secretase capacity to process its substrate, increase in saturation of γ-secretase with is substrate, and buildup of pathogenic interactions between different substrates and γ-secretase (Fig. 9) [7,29,65,69].

γ-Secretase is far from saturation with its substrate in physiological conditions in cells ([6,7,9,74], Fig. 9). For example, disease-causing APPsw mutation can increase enzyme activity by as much as 10 to 50 fold [44]! Such increase in γ-secretase activity is possible only if the enzyme is far below saturation in healthy conditions [17,75]. All enzymes in cells are far below saturation with their substrate [17,45,75,76]. Sub-saturated enzymes give the fastest and linear response to changes in metabolism [17,45,75,76]. Such conditions give the cells maximal control of the enzyme activity and metabolism [17,75,76] (Fig. 9). Sub-saturated enzymes can also favor metabolic regulation by supramolecular organization (Fig. 8 and [45,75,76]). We believe that measurements of the increase in the saturation of γ-secretase with its substrate, and related decrease in catalytic capacity of γ-secretase, can be new tools for the studies of the pathogenic changes in γ-secretase activity [7,29]. Such insights can be the basis for the development of the new early diagnostic tools [7,63,65]. The presented two-substrates-mechanism indicates that Alzheimer’s disease can be described as γ-secretase choking on its sticky substrates (Fig. 9).

### 3. C99-βCTF-APP substrate and different Aβ products can contribute to the pathogenic events

The presented two-substrate mechanism can also explain the apparently conflicting observations that both C99-βCTF-APP substrate and different Aβ products can lead to pathogenic events [36,65,69,77-79]. The general assumption is that amyloid proteins are released from γ-secretase to the lipid bilayer or even extracellular medium [3,78,80-82]. Then proteins start forming some kind of toxic Aβ oligomers by different mechanisms [81,83,84]. The spontaneous release of highly hydrophobic Aβ oligomers from hydrophobic lipid bilayer is not consistent with basic biophysical principles (supp. Fig. 3). The proposed secretion mechanisms have to be investigated [3,78].

We are proposing that the toxic events start when two substrates form contacts on the surface of γ-secretase (Figs. 3-5). The contacts can produce changes in γ-secretase structure (Fig. 7) that lead to an increase in Aβ (x-42)/ Aβ (x-40) ratio, accumulation of the longer more hydrophobic Aβ products, and accumulation of C99-βCTF-APP substrate [29]. Any of those three events can lead to interference with the physiological functions of γ-secretase, and thus, a large number of cytotoxic events [73,85,86]. The initial complex with C99-βCTF-APP and Aβ fragments can seed aggregation with other C99-βCTF-APP or Aβ proteins and ultimately lead to the formation of plaques [3,78,81]. The amyloid plaques are just the end result of the debris, that are gradually created in the saturation process (Fig. 9 [3,81]).

The presented two-substrate mechanism does not exclude the possibility that even a third C99-βCTF-APP, C83-αCTF-APP or Aβ molecule can bind to the presented two-substrate complex [9,10]. Such interactions could further facilitate the entire aggregation process [81]. The presented two-substrate mechanism does not exclude the possibility that an excess of C99-βCTF-APP substrate can cause toxic interfere with any other physiological functions of γ-secretase [1,2].

### 4. C99-βCTF-APP path is more likely to support the toxic two-substrate mechanism than C83-αCTF-APP path

The presented two-substrate-mechanism can explain why C99-βCTF-APP path is more toxic than C83-αCTF-APP path [3,81]. Both, C99-βCTF-APP or C83-αCTF-APP substrates, can form interactions with γ-secretase while it is catalytically processing any of its Aβ substrates (Fig. 3-5, and 7). The longer N-terminal domains in C99-βCTF-APP substrates have bigger interaction surfaces, and thus, have a bigger chance to form aggregates (Figs. 3-5). Thus, C99-βCTF-APP substrate and the corresponding Aβ oligomers, can produce bigger interference with the physiological functions of γ-secretase and lead to the related toxic steps [44].

### 5. The two-substrate mechanism and development of novel drug design strategies

Drug development studies were among the first to indicate that γ-secretase has two substrate-binding sites [8,9,12,16,30]. The presented two-substrate mechanism can be used for the optimization of future drug development strategies.

The first drug development strategy can be expansion of the future target list. A possible target can be any of the numerous physiological processes that control balance between γ-secretase activity and total amyloid metabolism (Fig. 9, [15,63,73,85,87]). The compounds that can decrease the catalytic capacity of γ-secretase can be used for the identification of different physiological processes that control γ-secretase activity and amyloid metabolism at pre-symptomatic stages (Fig. 9). Briefly, compounds such as semagacestat and avagacestat, can be used in healthy animals to gradually induce pathogenesis, by provoking gradual saturation of γ-secretase with its substrate (Fig. 9, [7,9]). The induced pathogenic events can be used for the description of the related physiological processes that control cellular levels of γ-secretase activity and/or total amyloid metabolism (Fig. 9).

The second drug-development strategy can be the development of competitive inhibitors of γ-secretase [17]. The competitive inhibitors should mimic the effects of the protective A673T mutation [17,41]. Attempts to design competitive inhibitors that target the active site tunnel have been challenging, and so far not successful [8,9,12,20]. Our results indicate two alternative strategies for the development of competitive inhibitors. First, the competitive inhibitors can be designed to facilitate the closing of the nicastrin ectodomain (Figs. 3-4 and 6). Second, the competitive inhibitors can be designed to bind to C99-βCTF-APP molecules and control its dimerization ([33,34,37], and Figs. 3-5 and 7). The development of compounds that target C99-βCTF-APP molecules is extremely difficult [81]. First, C99-βCTF-APP structure is highly dynamic and poorly defined target for effective drug-development efforts (Figs. 1-2, and [33,34,37]). Second, the prepared drugs have to compete with other molecules that bind to C99-βCTF-APP with high affinity ([37] and Fig. 1C).

The presented two-substrate mechanism can explain why the earlier studies which showed that multiple substrate and drug molecules can cooperate or compete in binding to γ-secretase [6,7,9,11,12]. Such competition can be affected by FAD mutations [6,7].

### Concluding remarks

The presented docking studies have been designed as an extension of earlier structural and docking studies [21-23,25,26,32]. However, the presented conclusions can be valid even if we do not know the precise substrate docking mechanism. Both, C99-βCTF-APP or C83-αCTF-APP substrates, can interact with γ-secretase while the enzyme is processing any of its Aβ substrates (Figs. 3-5 and 7, supp. videos 2 and 3). The interactions can be only transient contacts or a very specific complex. Any of those interactions can affect dynamic conformational changes that control processive catalysis by γ-secretase, but to a different extent ([25,32], supp. video 3). The number of different interactions presented in this study is limited by the introductory scope of this study, and additional mechanistic insights must be presented in future studies.

The presented mechanisms support proposals that the majority of uncertainties and inconstancies in studies of γ-secretase can be eliminated by controlling saturation of γ-secretase with its substrate [6,9,11-13,17,18,88]. All studies of γ-secretase activity can have a competition between substrates binding to γ-secretase (Fig. 3-5) and substrates binding to other substrates (Fig. 2). In cell-based studies dimerization of C99-βCTF-APP molecules is controlled by cell physiology, and could get out of control in cells that have unphysiological overexpression of C99-βCTF-APP molecules [6,88]. In enzyme-based studies dimerization between C99-βCTF-APP molecules can be minimized by starting the assays with the substrates that come immediately after elution from the affinity column by low pH [10]. The measurements at different saturations require extra efforts and costs. However, such measurements can give consistent results and sustained progress in enzyme-based, cell-based and computational studies [7-11,13,40,53,54,88].

This study showed that multiscale MD studies can be a crucial tool for studies of the molecular basis of Alzheimer’s disease and the related drug-development efforts [8,20-25]. The functional features of γ-secretase, C99-βCTF-APP, C83-αCTF-APP, and Aβ molecules depend on highly dynamic structures. Numerous transient contacts between flexible sites are driven by freely accessible charged, polar, and hydrophobic residues. The functions of such dynamic and sticky molecules cannot be captured by some of the structural studies [20,32,33,81].

## Materials and Methods

### Preparation of molecular structures for multiscale molecular dynamics (MD) calculations

All MD studies started with full-length C99-βCTF-APP structures that were built from fragments of available NMR conformers (Val13 to Tyr58, numeration based on PDB: 2LP1, [33]). The missing parts in NMR structures at the N-terminal end (residues 1 to 12) and the C-terminal end (residues 59 to 99) were built in several steps. First, the structures were prepared with no secondary structure presumptions using Modeller 9.17 [89], i.e. as fully extended forms (supp. video 1). Second, possible conformers in cholesterol-lipid-bilayer were calculated using multiscale MD studies and CHARMM-GUI tools [90]. Possible conformers were defined using coarse-grained MD studies that can depict as much as 20 microseconds of molecular events [28] (Fig. 1 and 2 A-C, supp. video 1) [90,91]. Finally, selected conformers and specific binding interactions, have been explored to atomic details by converting selected coarse-grained structures to all-atom structures and all-atom MD calculations (Fig. 2C) (AA-MD) [28,92]. Before different MD calculations, the transmembrane section was positioned in cholesterol-lipid-bilayer using OPM protocols [91].

Cryo-EM structures (PDB: 6IYC, [32]) were used to prepare γ-secretase structures with Aβ substrates of different lengths bound in the active site tunnel. The missing loops in γ-secretase structures and the missing N-terminal domain of the bound substrate were built with no secondary structure presumptions using Modeller 9.17 [89]. Modeller 9.17 can give between 5 and 10 conformers for short protein loops, which can be further optimized in multiscale MD protocols. Modeller 9.17 can give N-terminal domain of the bound substrate to different extent exposed or hidden by the nicastrin ectodomain. γ-Secretase structures with the substrate N-terminal domain exposed to different extent were used to study how the ectodomain can affect the enzyme functions. Cryo-EM structures (PDB: 6IYC, [32]) could not capture nicastrin ectodomain in closed conformation, what indicates that the function of closed ectodomain depends on multiple conformers. MD calculations used γ-secretase structures with the substrate-bound (PDB: 6IYC, [32], total 1355 residues), and γ-secretase structures without the bound substrate (PDB: 5FN2, [31], total 1309 residues). 6IYC and 5FN2 structures show differences between the structures with the active site tunnel in an open and closed conformation.

Structures of human BACE1 molecules (PDB: 4FGX, [46]) have been used with, and without bound inhibitor, to analyze functions of the active-site loops [45]. The missing 5 amino acid long loops have been prepared using Modeller 9.17 tool in UCSF Chimera [89].

Proteins were placed in a cholesterol-lipid bilayer that was designed to support catalytically relevant presenilin structures [93,94]. Main focus was on the active site Asp257 and Asp385 in presenilin 1 (Fig. 7 B-C, supp. Fig. 3 and [93]). The physiological relevance of the calculated presenilin structures was analyzed by comparing different mechanistic studies of the active site of aspartic protease ([25,95], supp. Fig. 3). We have also compared the calculated and experimental pKa for the active site aspartates [38,96]. Two different protocols were used to calculate pKa values. PropKa calculations for Asp257 is 6.9 and for Asp385 is and 6.8 [38]. Delphi calculations for Asp257 is 6.9 and for Asp385 is and 6.7 [97]. These values are comparable to the experimental pKa values [96]. We found that homogenous POPC bilayers that have been frequently used can produce some artifacts. Homogenous POPC membranes make an artificial charge distribution on the bilayer surface that can affect structures of γ-secretase and its substrate. POPC membranes have also a loose packing that cannot support catalytically optimal presenilin structures ([95], supp. Fig. 3). Studies of γ-secretase activity showed that a mixture of CHAPSO (cholesterol), POPE and POPC is crucial for the enzyme activity [10].

### Multiscale molecular dynamic studies of protein-protein interaction

Studies of protein-protein docking are an iterative process, a sequence of complementary steps [27,28]. All docking studies started with the two proteins placed in different rotations, about 5 to 30 Å away from each other. Different rotations and different distances were used to analyze to what extent complex formation, interaction sites, and the related conformations can be affected by the initial setup. The results from different calculations have been mutually compared and correlated with the available literature. The aim was to repeat docking studies multiple times with slightly different setup, with a desire to map differences and common features that can be observed from multiple events. Docking studies were gradually optimized with attempt to find conformers and contact sites that make maximal contact surfaces driven by free diffusion. The results of docking studies have been also correlated with different studies of γ–secretase activity.

All multiscale MD studies started with coarse-grained and steered MD calculations [27,98]. The aim was to explore a full range of possible molecular conformations, in search for conformers that could affect molecular interactions to a different degree. Selected structures from coarse-grained calculations were further subjected to a more detailed structural analysis at the level of individual atoms in all-atom MD studies [90,92,99].

In all docking studies with two free C99-β-CTF-APP molecules complex formation is at first driven by diffusion. Diffusion is greatly simplified, when both proteins have only mobility within two-dimensional lipid bilayer (supp. video 1). The diffusion distances between two molecules have a relatively small effect on the rate of complex formation (supp. video 1). The complex formation is primarily affected by the flexibility of C99-β-CTF-APP molecules, and by competition between intramolecular and intermolecular interactions. The relative orientation between the interacting molecules can affect the initial contact sites and the rate of interaction buildup. We focused our attention on the search for conformers that can give the biggest contact surface (supp. video 1).

All docking studies with free C99-β-CTF-APP and γ-secretase started with C99-β-CTF-APP structures positioned in different rotations facing TM2 and TM3 on presenilin subunit from distances between 5 to 30 Å [26,32]. TM2 and TM3 parts in the presenilin subunit have been frequently described as a part of a possible docking site [21,23,26]. Different distances and different rotations of C99-β-CTF-APP molecules have been used to explore the rate of interaction buildup, the number, and the position of interaction sites. Complex formation is greatly simplified when both proteins have mobility restricted within a two-dimensional lipid bilayer. We find in all docking studies that the rate of interaction buildup, the number, and the position of interaction sites, are affected by the rate of closing of nicastrin ectodomain and its different conformations. We focused attention on the search for conformers that support contacts between cytosolic end of presenilin 1 and C-terminal section of free C99-β-CTF-APP substrate (supp. video 3).

BACE1 molecules were not anchored to the membrane in docking studies [46]. Thus, with BACE1 molecules, the free diffusion to the surrounding solution is always in competition with the interaction with nicastrin ectodomain. The free diffusion makes docking studies with BACE1 more rigorous, relative to the membrane embedded studies, where proteins are trapped in lipid bilayers and have limited diffusion and rotation mobility. BACE1 needs to dock to nicastrin ectodomain in specific orientations. C-terminal domain of BACE1 has to be oriented towards the membrane surface, to satisfy the position of its transmembrane domain. BACE1 also needs to dock in the plane that is proximal to TM2 and TM3 on presenilin 1, to satisfy the criteria for substrate channeling [45]. We found that similar to docking interactions with free C99-β-CTF-APP molecules, the rate of interaction buildup, the number, and the position of interaction sites, are all affected by the rate of closing of nicastrin ectodomain and by its different conformations.

### Coarse-grained molecular dynamics calculations

Coarse-grained (CG) MD calculations with γ-secretase and/or its substrates used MARTINI 2.2 force field [98]. CHARMM-GUI protocol [99] was used to prepare simulation box with γ-secretase positioned in cholesterol-lipid-bilayer based on outputs from OPM protocols [91]. The smallest prepared box was 315 Å x 315 Å x 396 Å, mixed lipid bilayer have 1612 lipid molecules, 106668 water molecules, 1384 Na+ ions, 1278 Cl-ions in a box. Periodic boundary conditions employed in all directions first with NVT and second with NPT boundaries applied. The cholesterol-lipid-bilayer was assembled as phosphatidylcholine (POPC), 340 molecules (21%); phosphatidylethanolamine (POPE), 176 molecules (11%); phosphatidic acid (POPA), 16 molecules (1%); phosphatidylserine (POPS), 64 molecules (4%); sphingomyelin (PSM), 96 molecules (6%); phosphatidylinositol (POPI), 32 molecules (2%); cholesterol (CHOL), 880 molecules (55%).

The prepared simulation box with the protein in the bilayer has been subjected to two rounds of minimization, with the integrator set to steep, and the number of integration steps set to 5000, until default was reached. Four rounds of equilibration followed, with the integrator set to md, and with gradually increasing the time step from 5, 10, 15, 20 femtoseconds. For all minimization and equilibration steps pressure coupling was set to Berendsen, semiisotropic, with tau_p set to 5.0, and compressibility set to = 3e-4. Cutoff-scheme was set to Verlet, ns_type was set to grid, with verlet-buffer-tolerance = 0.005, and epsilon_r set to 15. Coulombtype was setup to reaction-field, rcoulomb = 1.1, vdw_type= cutoff, vdw-modifier, Potential-shift-verlet, rvdw= 1.1. Tcoupl = v-rescale, tc-grps = protein membrane solute, tau_t = 1.0 1.0 1.0, ref_t = 303.15 K.

The MD calculations used between 0.5 to 1 billion integration steps, with the integration time set to 20 femtoseconds. The calculation results were recorded in 1000 to 2000 frames, to depict in a total of 10 to 20 microseconds of molecular events. The pressure (1 atm) and temperature (300 K) were held constant using Langevin thermostat with a collision frequency of 1 ps-1. Bonds with hydrogen atoms were constrained using the SHAKE algorithm, while the long-range electrostatic interactions have been calculated using the Particle Mesh Ewald method. The calculations used between 20 to 40 nodes on Atos, Bullx DLC 720 system, and took about 3 to 5 days. Each node had two Xeon E5-2690 12C 2.6GHz (24 physical cores per node) and 64 GB RAM.

### All atom molecular dynamics calculations

All atom molecular dynamics calculations (AA-MD) with γ-secretase and/or its substrates positioned in cholesterol-lipid-bilayer have been prepared using CHARMM-GUI Membrane Builder with CHARMM36a force field [90,92]. Proteins were positioned in bilayer following OPM protocols [91]. OPM structures were placed into, typical water box had 1355 residues, 708 lipid molecules, 148692 TIP3 water molecules, 419 Na+ ions and 414 Cl-ions in a 153 Å x 153 Å x 247 Å box (150 mM NaCl). The cholesterol-lipid-bilayer was prepared as: phosphatidylcholine (POPC), 152 molecules (21%); phosphatidylethanolamine (POPE), 78 molecules (11%); phosphatidic acid (POPA), 8 molecules (1%); phosphatidylserine (POPS), 28 molecules (4%); sphingomyelin (PSM), 42 molecules (6%); phosphatidylinositol (POPI), 14 molecules (2%); cholesterol (CHOL) 386 molecules (55%).

The prepared simulation box was subjected to minimization, integrator = steep, emtol = 1000.0, nsteps = 5000, nstlist= 10, cutoff-scheme = Verlet, rlist= 1.2, vdwtype = Cut-off, vdw-modifier = Force-switch, rvdw_switch = 1.0, rvdw= 1.2, coulombtype = pme, rcoulomb = 1.2, minimsed in 5000 steps. The system was subsequently relaxed in 6 equilibration steps with the gradually increasing integration time. The setup for the equilibration steps was: integrator = md, cutoff-scheme = Verlet, nstlist= 20, rlist= 1.2, coulombtype = pme, rcoulomb = 1.2, vdwtype = Cut-off, vdw-modifier = Force-switch, rvdw_switch = 1.0, rvdw = 1.2, tcoupl = berendsen, tc_grps = PROT MEMB SOL_ION, tau_t = 1.0, ref_t = 303.15, pcoupl = berendsen, pcoupltype = semiisotropic, tau_p= 5.0, compressibility= 4.5e-5 4.5e-5, ref_p= 1.0, constraints= h-bonds, constraint_algorithm= LINCS, continuation= yes, comm_grps= PROT MEMB SOL_ION, refcoord_scaling= com.

MD calculations took 3 to 6 days on Atos, Bullx DLC 720 system, using 20 to 50 nodes. Each node had two Xeon E5-2690v3 12C 2.6GHz each processors with 24 physical cores. MD calculation used system with the temperature set to 303.15 K, Nose-Hoover coupling, and the pressure set to 1.0 bar using semi-isotropic Parinello-Rahman coupling. Calculations took between 100 to 200 million steps, with the step size set to 2 femtoseconds. The results were recorded in 150 to 200 frames, to depict between 200 to 400 nanoseconds of molecular events.

### Statistical analysis of molecular dynamics results

Molecular trajectories and conformations calculated in molecular dynamics studies have been analyzed quantitatively using statistical protocols from Bio3D package and program R 3.6.2 [39]. Calculations of root-mean-square deviation (RMSD) values can be used to monitor convergence in the extent of molecular mobility. Calculations of root-mean-square fluctuation (RMSF) can be used for identifications of difference in mobility at the specific molecular sites. In extended forms RMSF values can be used to compare results from different MD calculations that used the same setup.

## Supporting information

supplemental file description

## Author Contributions

Conceptualization, Ž.M.S.; methodology, Ž.M.S., L.O.; software, Ž.M.S., L.O.; validation, Ž.M.S., L.O.; formal analysis, Ž.M.S., L.O.; investigation, Ž.M.S., L.O.; resources, Ž.M.S. V.Š.J; data curation, Ž.M.S. V.Š.J; writing—original draft preparation, Ž.M.S., L.O.; writing—review and editing, Ž.M.S.; visualization, Ž.M.S., L.O.; supervision, Ž.M.S., V.Š.J; project administration, Ž.M.S., V.J.Š.; funding acquisition, Ž.M.S., V.J.Š., All authors have read and agreed to the published version of the manuscript.

## Acknowledgments

Ž.M.S. was a recipient of funds from the Croatian Science Foundation’s project number O-1505-2015, and was—for a period of time—employed in medical biochemistry laboratory at the Psychiatric Hospital Rab. The authors gratefully acknowledge the services of Gordan Janeš and Draško Tomić who have provided crucial computational expertise as a part of the Center for Advanced Computing and Modeling. We apologize that we could not include many of the relevant citations due to space limitations.

## References

1. Scheltens, P.; Blennow, K.; Breteler, M.M.; de Strooper, B.; Frisoni, G.B.; Salloway, S.; Van der Flier, W.M. Alzheimer’s disease. Lancet (London, England) 2016, 388, 505–517, doi:10.1016/s0140-6736(15)01124-1.

2. Imbimbo, B.P.; Panza, F.; Frisardi, V.; Solfrizzi, V.; D’Onofrio, G.; Logroscino, G.; Seripa, D.; Pilotto, A. Therapeutic intervention for Alzheimer’s disease with gamma-secretase inhibitors: still a viable option? Expert Opin Investig Drugs 2010, 20, 325–341.

3. Castro, M.A.; Hadziselimovic, A.; Sanders, C.R. The vexing complexity of the amyloidogenic pathway. Protein science : a publication of the Protein Society 2019, 28, 1177–1193, doi:10.1002/pro.3606.

4. Toyn, J.H.; Ahlijanian, M.K. Interpreting Alzheimer’s disease clinical trials in light of the effects on amyloid-β. Alzheimers Res Ther 2014, 6, 14, doi:10.1186/alzrt244.

5. Sambamurti, K.; Greig, N.H.; Utsuki, T.; Barnwell, E.L.; Sharma, E.; Mazell, C.; Bhat, N.R.; Kindy, M.S.; Lahiri, D.K.; Pappolla, M.A. Targets for AD treatment: conflicting messages from gamma-secretase inhibitors. J Neurochem 2011, 117, 359–374.

6. Burton, C.R.; Meredith, J.E.; Barten, D.M.; Goldstein, M.E.; Krause, C.M.; Kieras, C.J.; Sisk, L.; Iben, L.G.; Polson, C.; Thompson, M.W., et al. The amyloid-beta rise and gamma-secretase inhibitor potency depend on the level of substrate expression. The Journal of biological chemistry 2008, 283, 22992–23003.

7. Svedružić Ž, M.; Popović, K.; Šendula-Jengić, V. Decrease in catalytic capacity of γ-secretase can facilitate pathogenesis in sporadic and Familial Alzheimer’s disease. Mol Cell Neurosci 2015, 67, 55–65, doi:10.1016/j.mcn.2015.06.002.

8. Svedružić Ž, M.; Vrbnjak, K.; Martinović, M.; Miletić, V. Structural Analysis of the Simultaneous Activation and Inhibition of γ-Secretase Activity in the Development of Drugs for Alzheimer’s Disease. Pharmaceutics 2021, 13, doi:10.3390/pharmaceutics13040514.

9. Svedružić, Z.M.; Popovic, K.; Sendula-Jengic, V. Modulators of gamma-secretase activity can facilitate the toxic side-effects and pathogenesis of Alzheimer’s disease. PloS one 2013, 8, e50759, doi:10.1371/journal.pone.0050759.

10. Svedružić, Z.M.; Popovic, K.; Smoljan, I.; Sendula-Jengic, V. Modulation of gamma-Secretase Activity by Multiple Enzyme-Substrate Interactions: Implications in Pathogenesis of Alzheimer’s Disease. PloS one 2012, 7, e32293.

11. Yagishita, S.; Morishima-Kawashima, M.; Tanimura, Y.; Ishiura, S.; Ihara, Y. DAPT-induced intracellular accumulations of longer amyloid beta-proteins: further implications for the mechanism of intramembrane cleavage by gamma-secretase. Biochemistry 2006, 45, 3952–3960.

12. Walsh, R. Are improper kinetic models hampering drug development? PeerJ 2014, 2, e649, doi:10.7717/peerj.649.

13. Kakuda, N.; Funamoto, S.; Yagishita, S.; Takami, M.; Osawa, S.; Dohmae, N.; Ihara, Y. Equimolar production of amyloid beta-protein and amyloid precursor protein intracellular domain from beta-carboxyl-terminal fragment by gamma-secretase. The Journal of biological chemistry 2006, 281, 14776–14786.

14. Yin, Y.I.; Bassit, B.; Zhu, L.; Yang, X.; Wang, C.; Li, Y.M. {gamma}-Secretase Substrate Concentration Modulates the Abeta42/Abeta40 Ratio: implications for Alzheimer’s disease. The Journal of biological chemistry 2007, 282, 23639–23644.

15. Hur, J.Y.; Frost, G.R.; Wu, X.; Crump, C.; Pan, S.J.; Wong, E.; Barros, M.; Li, T.; Nie, P.; Zhai, Y., et al. The innate immunity protein IFITM3 modulates γ-secretase in Alzheimer’s disease. Nature 2020, 586, 735–740, doi:10.1038/s41586-020-2681-2.

16. Wolfe, M.S. Probing Mechanisms and Therapeutic Potential of γ-Secretase in Alzheimer’s Disease. Molecules (Basel, Switzerland) 2021, 26, doi:10.3390/molecules26020388.

17. Fersht, A. Structure and Mechanism in Protein Science: A Guide to Enzyme Catalysis and Protein Folding, 4th ed.; World Scientific Publishing Co: 2018; pp. 656

18. Motulsky, H.; Christopoulos, A. Fitting Models to Biological Data Using Linear and Nonlinear Regression: A Practical Guide to Curve Fitting Oxford University Press, USA; 1 edition 2004; pp. 352.

19. Johnson, K.A. Fitting enzyme kinetic data with KinTek global kinetic explorer. Methods in enzymology 2009, 467, 601–626.

20. Yang, G.; Zhou, R.; Guo, X.; Yan, C.; Lei, J.; Shi, Y. Structural basis of γ-secretase inhibition and modulation by small molecule drugs. Cell 2021, 184, 521–533 e514, doi:10.1016/j.cell.2020.11.049.

21. Aguayo-Ortiz, R.; Chávez-García, C.; Straub, J.E.; Dominguez, L. Characterizing the structural ensemble of γ-secretase using a multiscale molecular dynamics approach. Chem Sci 2017, 8, 5576–5584, doi:10.1039/c7sc00980a.

22. Bolduc, D.M.; Montagna, D.R.; Gu, Y.; Selkoe, D.J.; Wolfe, M.S. Nicastrin functions to sterically hinder γ-secretase-substrate interactions driven by substrate transmembrane domain. Proceedings of the National Academy of Sciences of the United States of America 2016, 113, E509–518, doi:10.1073/pnas.1512952113.

23. Lee, J.Y.; Feng, Z.; Xie, X.Q.; Bahar, I. Allosteric Modulation of Intact γ-Secretase Structural Dynamics. Biophys J 2017, 113, 2634–2649, doi:10.1016/j.bpj.2017.10.012.

24. Pantelopulos, G.A.; Straub, J.E.; Thirumalai, D.; Sugita, Y. Structure of APP-C99(1-99) and implications for role of extra-membrane domains in function and oligomerization. Biochimica et biophysica acta. Biomembranes 2018, 1860, 1698–1708, doi:10.1016/j.bbamem.2018.04.002.

25. Bhattarai, A.; Devkota, S.; Do, H.N.; Wang, J.; Bhattarai, S.; Wolfe, M.S.; Miao, Y. Mechanism of Tripeptide Trimming of Amyloid β-Peptide 49 by γ-Secretase. Journal of the American Chemical Society 2022, 10.1021/jacs.1c10533, doi:10.1021/jacs.1c10533.

26. Bai, X.C.; Rajendra, E.; Yang, G.; Shi, Y.; Scheres, S.H. Sampling the conformational space of the catalytic subunit of human γ-secretase. Elife 2015, 4, doi:10.7554/eLife.11182.

27. Van Der Spoel, D.; Lindahl, E.; Hess, B.; Groenhof, G.; Mark, A.E.; Berendsen, H.J. GROMACS: fast, flexible, and free. Journal of computational chemistry 2005, 26, 1701–1718.

28. Roel-Touris, J.; Bonvin, A. Coarse-grained (hybrid) integrative modeling of biomolecular interactions. Computational and structural biotechnology journal 2020, 18, 1182–1190, doi:10.1016/j.csbj.2020.05.002.

29. Checler, F.; Afram, E.; Pardossi-Piquard, R.; Lauritzen, I. Is γ-secretase a beneficial inactivating enzyme of the toxic APP C-terminal fragment C99? The Journal of biological chemistry 2021, 296, 100489, doi:10.1016/j.jbc.2021.100489.

30. Kornilova, A.Y.; Bihel, F.; Das, C.; Wolfe, M.S. The initial substrate-binding site of gamma-secretase is located on presenilin near the active site. Proceedings of the National Academy of Sciences of the United States of America 2005, 102, 3230–3235.

31. Bai, X.C.; Yan, C.; Yang, G.; Lu, P.; Ma, D.; Sun, L.; Zhou, R.; Scheres, S.H.W.; Shi, Y. An atomic structure of human γ-secretase. Nature 2015, 525, 212–217, doi:10.1038/nature14892.

32. Zhou, R.; Yang, G.; Guo, X.; Zhou, Q.; Lei, J.; Shi, Y. Recognition of the amyloid precursor protein by human γ-secretase. Science 2019, 363, doi:10.1126/science.aaw0930.

33. Barrett, P.J.; Song, Y.; Van Horn, W.D.; Hustedt, E.J.; Schafer, J.M.; Hadziselimovic, A.; Beel, A.J.; Sanders, C.R. The amyloid precursor protein has a flexible transmembrane domain and binds cholesterol. Science 2012, 336, 1168–1171.

34. Richter, L.; Munter, L.M.; Ness, J.; Hildebrand, P.W.; Dasari, M.; Unterreitmeier, S.; Bulic, B.; Beyermann, M.; Gust, R.; Reif, B., et al. Amyloid beta 42 peptide (Abeta42)-lowering compounds directly bind to Abeta and interfere with amyloid precursor protein (APP) transmembrane dimerization. Proceedings of the National Academy of Sciences of the United States of America 2010, 107, 14597–14602.

35. Eggert, S.; Midthune, B.; Cottrell, B.; Koo, E.H. Induced dimerization of the amyloid precursor protein leads to decreased amyloid-beta protein production. The Journal of biological chemistry 2009, 284, 28943–28952.

36. Gorman, P.M.; Kim, S.; Guo, M.; Melnyk, R.A.; McLaurin, J.; Fraser, P.E.; Bowie, J.U.; Chakrabartty, A. Dimerization of the transmembrane domain of amyloid precursor proteins and familial Alzheimer’s disease mutants. BMC Neurosci 2008, 9, 17.

37. Song, Y.; Hustedt, E.J.; Brandon, S.; Sanders, C.R. Competition between homodimerization and cholesterol binding to the C99 domain of the amyloid precursor protein. Biochemistry 2013, 52, 5051–5064, doi:10.1021/bi400735x.

38. Li, H.; Robertson, A.D.; Jensen, J.H. Very fast empirical prediction and rationalization of protein pKa values. Proteins 2005, 61, 704–721, doi:10.1002/prot.20660.

39. Grant, B.J.; Skjærven, L.; Yao, X.Q. Comparative Protein Structure Analysis with Bio3D-Web. Methods Mol Biol 2020, 2112, 15–28, doi:10.1007/978-1-0716-0270-6_2.

40. Kakuda, N.; Takami, M.; Okochi, M.; Kasuga, K.; Ihara, Y.; Ikeuchi, T. Switched Aβ43 generation in familial Alzheimer’s disease with presenilin 1 mutation. Translational psychiatry 2021, 11, 558, doi:10.1038/s41398-021-01684-1.

41. Jonsson, T.; Atwal, J.K.; Steinberg, S.; Snaedal, J.; Jonsson, P.V.; Bjornsson, S.; Stefansson, H.; Sulem, P.; Gudbjartsson, D.; Maloney, J., et al. A mutation in APP protects against Alzheimer’s disease and age-related cognitive decline. Nature 2012, 488, 96–99.

42. Lu, P.; Bai, X.C.; Ma, D.; Xie, T.; Yan, C.; Sun, L.; Yang, G.; Zhao, Y.; Zhou, R.; Scheres, S.H.W., et al. Three-dimensional structure of human γ-secretase. Nature 2014, 512, 166–170, doi:10.1038/nature13567.

43. Liu, L.; Ding, L.; Rovere, M.; Wolfe, M.S.; Selkoe, D.J. A cellular complex of BACE1 and γ-secretase sequentially generates Aβ from its full-length precursor. The Journal of cell biology 2019, 218, 644–663, doi:10.1083/jcb.201806205.

44. McDade, E.; Voytyuk, I.; Aisen, P.; Bateman, R.J.; Carrillo, M.C.; De Strooper, B.; Haass, C.; Reiman, E.M.; Sperling, R.; Tariot, P.N., et al. The case for low-level BACE1 inhibition for the prevention of Alzheimer disease. Nature reviews. Neurology 2021, 17, 703–714, doi:10.1038/s41582-021-00545-1.

45. Svedružić Ž, M.; Odorčić, I.; Chang, C.H.; Svedružić, D. Substrate Channeling via a Transient Protein-Protein Complex: The case of D-Glyceraldehyde-3-Phosphate Dehydrogenase and L-Lactate Dehydrogenase. Sci Rep 2020, 10, 10404, doi:10.1038/s41598-020-67079-2.

46. Liu, Y.; Zhang, W.; Li, L.; Salvador, L.A.; Chen, T.; Chen, W.; Felsenstein, K.M.; Ladd, T.B.; Price, A.R.; Golde, T.E., et al. Cyanobacterial peptides as a prototype for the design of potent β-secretase inhibitors and the development of selective chemical probes for other aspartic proteases. Journal of medicinal chemistry 2012, 55, 10749–10765, doi:10.1021/jm301630s.

47. Bhattarai, S.; Devkota, S.; Meneely, K.M.; Xing, M.; Douglas, J.T.; Wolfe, M.S. Design of Substrate Transmembrane Mimetics as Structural Probes for γ-Secretase. Journal of the American Chemical Society 2020, 142, 3351–3355, doi:10.1021/jacs.9b13405.

48. Dehury, B.; Tang, N.; Kepp, K.P. Molecular dynamics of C99-bound γ-secretase reveal two binding modes with distinct compactness, stability, and active-site retention: implications for Aβ production. Biochem J 2019, 476, 1173–1189, doi:10.1042/bcj20190023.

49. Shen, J.; Kelleher, R.J., 3rd. The presenilin hypothesis of Alzheimer’s disease: evidence for a loss-of-function pathogenic mechanism. Proceedings of the National Academy of Sciences of the United States of America 2007, 104, 403–409, doi:10.1073/pnas.0608332104.

50. Wang, S.C.; Oelze, B.; Schumacher, A. Age-specific epigenetic drift in late-onset Alzheimer’s disease. PloS one 2008, 3, e2698, doi:10.1371/journal.pone.0002698.

51. Ganguly, G.; Chakrabarti, S.; Chatterjee, U.; Saso, L. Proteinopathy, oxidative stress and mitochondrial dysfunction: cross talk in Alzheimer’s disease and Parkinson’s disease. Drug design, development and therapy 2017, 11, 797–810, doi:10.2147/dddt.s130514.

52. Hunter, S.; Arendt, T.; Brayne, C. The senescence hypothesis of disease progression in Alzheimer disease: an integrated matrix of disease pathways for FAD and SAD. Mol Neurobiol 2013, 48, 556–570, doi:10.1007/s12035-013-8445-3.

53. Qi, Y.; Morishima-Kawashima, M.; Sato, T.; Mitsumori, R.; Ihara, Y. Distinct mechanisms by mutant presenilin 1 and 2 leading to increased intracellular levels of amyloid beta-protein 42 in Chinese hamster ovary cells. Biochemistry 2003, 42, 1042–1052.

54. Qi-Takahara, Y.; Morishima-Kawashima, M.; Tanimura, Y.; Dolios, G.; Hirotani, N.; Horikoshi, Y.; Kametani, F.; Maeda, M.; Saido, T.C.; Wang, R., et al. Longer forms of amyloid beta protein: implications for the mechanism of intramembrane cleavage by gamma-secretase. The Journal of neuroscience : the official journal of the Society for Neuroscience 2005, 25, 436–445.

55. Wolfe, M.S. When loss is gain: reduced presenilin proteolytic function leads to increased Abeta42/Abeta40. Talking Point on the role of presenilin mutations in Alzheimer disease. EMBO reports 2007, 8, 136–140, doi:10.1038/sj.embor.7400896.

56. Cai, X.D.; Golde, T.E.; Younkin, S.G. Release of excess amyloid beta protein from a mutant amyloid beta protein precursor. Science 1993, 259, 514–516.

57. Citron, M.; Oltersdorf, T.; Haass, C.; McConlogue, L.; Hung, A.Y.; Seubert, P.; Vigo-Pelfrey, C.; Lieberburg, I.; Selkoe, D.J. Mutation of the beta-amyloid precursor protein in familial Alzheimer’s disease increases beta-protein production. Nature 1992, 360, 672–674.

58. Sleegers, K.; Brouwers, N.; Gijselinck, I.; Theuns, J.; Goossens, D.; Wauters, J.; Del-Favero, J.; Cruts, M.; van Duijn, C.M.; Van Broeckhoven, C. APP duplication is sufficient to cause early onset Alzheimer’s dementia with cerebral amyloid angiopathy. Brain : a journal of neurology 2006, 129, 2977–2983, doi:10.1093/brain/awl203.

59. Saura, C.A.; Choi, S.Y.; Beglopoulos, V.; Malkani, S.; Zhang, D.; Shankaranarayana Rao, B.S.; Chattarji, S.; Kelleher, R.J., 3rd; Kandel, E.R.; Duff, K., et al. Loss of presenilin function causes impairments of memory and synaptic plasticity followed by age-dependent neurodegeneration. Neuron 2004, 42, 23–36, doi:10.1016/s0896-6273(04)00182-5.

60. Guix, F.X.; Wahle, T.; Vennekens, K.; Snellinx, A.; Chavez-Gutierrez, L.; Ill-Raga, G.; Ramos-Fernandez, E.; Guardia-Laguarta, C.; Lleo, A.; Arimon, M., et al. Modification of gamma-secretase by nitrosative stress links neuronal ageing to sporadic Alzheimer’s disease. EMBO Mol Med 2012, 4, 660–673.

61. Refolo, L.M.; Eckman, C.; Prada, C.M.; Yager, D.; Sambamurti, K.; Mehta, N.; Hardy, J.; Younkin, S.G. Antisense-induced reduction of presenilin 1 expression selectively increases the production of amyloid beta42 in transfected cells. J Neurochem 1999, 73, 2383–2388.

62. Andreoli, V.; Trecroci, F.; La Russa, A.; Cittadella, R.; Liguori, M.; Spadafora, P.; Caracciolo, M.; Di Palma, G.; Colica, C.; Gambardella, A., et al. Presenilin enhancer-2 gene: identification of a novel promoter mutation in a patient with early-onset familial Alzheimer’s disease. Alzheimer’s & dementia : the journal of the Alzheimer’s Association 2011, 7, 574–578, doi:10.1016/j.jalz.2011.02.010.

63. Theuns, J.; Remacle, J.; Killick, R.; Corsmit, E.; Vennekens, K.; Huylebroeck, D.; Cruts, M.; Van Broeckhoven, C. Alzheimer-associated C allele of the promoter polymorphism -22C>T causes a critical neuron-specific decrease of presenilin 1 expression. Hum Mol Genet 2003, 12, 869–877.

64. Nishimura, M.; Nakamura, S.I.; Kimura, N.; Liu, L.; Suzuki, T.; Tooyama, I. Age-related modulation of gamma-secretase activity in non-human primate brains. J Neurochem 2012.

65. Tamayev, R.; D’Adamio, L. Inhibition of gamma-secretase worsens memory deficits in a genetically congruous mouse model of Danish dementia. Mol Neurodegener 2012, 7, 19.

66. Fukumoto, H.; Rosene, D.L.; Moss, M.B.; Raju, S.; Hyman, B.T.; Irizarry, M.C. Beta-secretase activity increases with aging in human, monkey, and mouse brain. Am J Pathol 2004, 164, 719–725.

67. Li, R.; Lindholm, K.; Yang, L.B.; Yue, X.; Citron, M.; Yan, R.; Beach, T.; Sue, L.; Sabbagh, M.; Cai, H., et al. Amyloid beta peptide load is correlated with increased beta-secretase activity in sporadic Alzheimer’s disease patients. Proceedings of the National Academy of Sciences of the United States of America 2004, 101, 3632–3637.

68. Rovelet-Lecrux, A.; Hannequin, D.; Raux, G.; Le Meur, N.; Laquerriere, A.; Vital, A.; Dumanchin, C.; Feuillette, S.; Brice, A.; Vercelletto, M., et al. APP locus duplication causes autosomal dominant early-onset Alzheimer disease with cerebral amyloid angiopathy. Nat Genet 2006, 38, 24–26.

69. Bourgeois, A.; Lauritzen, I.; Lorivel, T.; Bauer, C.; Checler, F.; Pardossi-Piquard, R. Intraneuronal accumulation of C99 contributes to synaptic alterations, apathy-like behavior, and spatial learning deficits in 3×TgAD and 2×TgAD mice. Neurobiology of aging 2018, 71, 21–31, doi:10.1016/j.neurobiolaging.2018.06.038.

70. Kern, A.; Behl, C. The unsolved relationship of brain aging and late-onset Alzheimer disease. Biochimica et biophysica acta 2009, 1790, 1124–1132, doi:10.1016/j.bbagen.2009.07.016.

71. Miners, J.S.; Jones, R.; Love, S. Differential changes in Aβ42 and Aβ40 with age. Journal of Alzheimer’s disease : JAD 2014, 40, 727–735, doi:10.3233/jad-132339.

72. Berchtold, N.C.; Cribbs, D.H.; Coleman, P.D.; Rogers, J.; Head, E.; Kim, R.; Beach, T.; Miller, C.; Troncoso, J.; Trojanowski, J.Q., et al. Gene expression changes in the course of normal brain aging are sexually dimorphic. Proceedings of the National Academy of Sciences of the United States of America 2008, 105, 15605–15610, doi:10.1073/pnas.0806883105.

73. Thathiah, A.; De Strooper, B. The role of G protein-coupled receptors in the pathology of Alzheimer’s disease. Nat Rev Neurosci 2011, 12, 73–87.

74. Jämsä, A.; Belda, O.; Edlund, M.; Lindström, E. BACE-1 inhibition prevents the γ-secretase inhibitor evoked Aβ rise in human neuroblastoma SH-SY5Y cells. J Biomed Sci 2011, 18, 76, doi:10.1186/1423-0127-18-76.

75. De la Fuente, I.M.; Martínez, L.; Carrasco-Pujante, J.; Fedetz, M.; López, J.I.; Malaina, I. Self-Organization and Information Processing: From Basic Enzymatic Activities to Complex Adaptive Cellular Behavior. Frontiers in genetics 2021, 12, 644615, doi:10.3389/fgene.2021.644615.

76. Van Noorden, C.J.; Jonges, G.N. Analysis of enzyme reactions in situ. Histochem J 1995, 27, 101–118.

77. Lauritzen, I.; Pardossi-Piquard, R.; Bauer, C.; Brigham, E.; Abraham, J.D.; Ranaldi, S.; Fraser, P.; St-George-Hyslop, P.; Le Thuc, O.; Espin, V., et al. The β-secretase-derived C-terminal fragment of βAPP, C99, but not Aβ, is a key contributor to early intraneuronal lesions in triple-transgenic mouse hippocampus. The Journal of neuroscience : the official journal of the Society for Neuroscience 2012, 32, 16243–11655a, doi:10.1523/jneurosci.2775-12.2012.

78. Mondragón-Rodríguez, S.; Gu, N.; Manseau, F.; Williams, S. Alzheimer’s Transgenic Model Is Characterized by Very Early Brain Network Alterations and β-CTF Fragment Accumulation: Reversal by β-Secretase Inhibition. Frontiers in cellular neuroscience 2018, 12, 121, doi:10.3389/fncel.2018.00121.

79. Gralle, M.; Botelho, M.G.; Wouters, F.S. Neuroprotective secreted amyloid precursor protein acts by disrupting amyloid precursor protein dimers. The Journal of biological chemistry 2009, 284, 15016–15025.

80. Gomes, G.N.; Levine, Z.A. Defining the Neuropathological Aggresome across in Silico, in Vitro, and ex Vivo Experiments. The journal of physical chemistry. B 2021, 125, 1974–1996, doi:10.1021/acs.jpcb.0c09193.

81. Chen, G.F.; Xu, T.H.; Yan, Y.; Zhou, Y.R.; Jiang, Y.; Melcher, K.; Xu, H.E. Amyloid beta: structure, biology and structure-based therapeutic development. Acta pharmacologica Sinica 2017, 38, 1205–1235, doi:10.1038/aps.2017.28.

82. Fan, J.; Donkin, J.; Wellington, C. Greasing the wheels of Abeta clearance in Alzheimer’s disease: the role of lipids and apolipoprotein E. Biofactors 2009, 35, 239–248.

83. Lauritzen, I.; Pardossi-Piquard, R.; Bourgeois, A.; Pagnotta, S.; Biferi, M.G.; Barkats, M.; Lacor, P.; Klein, W.; Bauer, C.; Checler, F. Intraneuronal aggregation of the β-CTF fragment of APP (C99) induces Aβ-independent lysosomal-autophagic pathology. Acta neuropathologica 2016, 132, 257–276, doi:10.1007/s00401-016-1577-6.

84. Axelsen, P.H.; Komatsu, H.; Murray, I.V. Oxidative stress and cell membranes in the pathogenesis of Alzheimer’s disease. Physiology (Bethesda) 2011, 26, 54–69.

85. Jurisch-Yaksi, N.; Sannerud, R.; Annaert, W. A fast growing spectrum of biological functions of γ-secretase in development and disease. Biochimica et biophysica acta 2013, 1828, 2815–2827, doi:10.1016/j.bbamem.2013.04.016.

86. Hunter, S.; Brayne, C. Integrating the molecular and the population approaches to dementia research to help guide the future development of appropriate therapeutics. Biochem Pharmacol 2014, 88, 652–660, doi:10.1016/j.bcp.2013.12.015.

87. Kisby, B.; Jarrell, J.T.; Agar, M.E.; Cohen, D.S.; Rosin, E.R.; Cahill, C.M.; Rogers, J.T.; Huang, X. Alzheimer’s Disease and Its Potential Alternative Therapeutics. J Alzheimers Dis Parkinsonism 2019, 9, doi:10.4172/2161-0460.1000477.

88. Armbrust, F.; Bickenbach, K.; Marengo, L.; Pietrzik, C.; Becker-Pauly, C. The Swedish dilemma -the almost exclusive use of APPswe-based mouse models impedes adequate evaluation of alternative β-secretases. Biochimica et biophysica acta. Molecular cell research 2022, 1869, 119164, doi:10.1016/j.bbamcr.2021.119164.

89. Webb, B.; Sali, A. Comparative Protein Structure Modeling Using MODELLER. Current protocols in bioinformatics 2016, 54, 5 6 1–5 6 37, doi:10.1002/cpbi.3.

90. Lee, J.; Patel, D.S.; Ståhle, J.; Park, S.J.; Kern, N.R.; Kim, S.; Lee, J.; Cheng, X.; Valvano, M.A.; Holst, O., et al. CHARMM-GUI Membrane Builder for Complex Biological Membrane Simulations with Glycolipids and Lipoglycans. J Chem Theory Comput 2019, 15, 775–786, doi:10.1021/acs.jctc.8b01066.

91. Lomize, M.A.; Pogozheva, I.D.; Joo, H.; Mosberg, H.I.; Lomize, A.L. OPM database and PPM web server: resources for positioning of proteins in membranes. Nucleic Acids Res 2012, 40, D370–376, doi:10.1093/nar/gkr703.

92. Lee, J.; Cheng, X.; Swails, J.M.; Yeom, M.S.; Eastman, P.K.; Lemkul, J.A.; Wei, S.; Buckner, J.; Jeong, J.C.; Qi, Y., et al. CHARMM-GUI Input Generator for NAMD, GROMACS, AMBER, OpenMM, and CHARMM/OpenMM Simulations Using the CHARMM36 Additive Force Field. J Chem Theory Comput 2016, 12, 405–413, doi:10.1021/acs.jctc.5b00935.

93. Audagnotto, M.; Kengo Lorkowski, A.; Dal Peraro, M. Recruitment of the amyloid precursor protein by γ-secretase at the synaptic plasma membrane. Biochem Biophys Res Commun 2018, 498, 334–341, doi:10.1016/j.bbrc.2017.10.164.

94. Aguayo-Ortiz, R.; Straub, J.E.; Dominguez, L. Influence of membrane lipid composition on the structure and activity of γ-secretase. Physical chemistry chemical physics : PCCP 2018, 20, 27294–27304, doi:10.1039/c8cp04138e.

95. Krzemińska, A.; Moliner, V.; Świderek, K. Dynamic and Electrostatic Effects on the Reaction Catalyzed by HIV-1 Protease. Journal of the American Chemical Society 2016, 138, 16283–16298, doi:10.1021/jacs.6b06856.

96. Li, Y.M.; Lai, M.T.; Xu, M.; Huang, Q.; DiMuzio-Mower, J.; Sardana, M.K.; Shi, X.P.; Yin, K.C.; Shafer, J.A.; Gardell, S.J. Presenilin 1 is linked with gamma-secretase activity in the detergent solubilized state. Proceedings of the National Academy of Sciences of the United States of America 2000, 97, 6138–6143.

97. Pahari, S.; Sun, L.; Basu, S.; Alexov, E. DelPhiPKa: Including salt in the calculations and enabling polar residues to titrate. Proteins 2018, 86, 1277–1283, doi:10.1002/prot.25608.

98. Arnarez, C.; Uusitalo, J.J.; Masman, M.F.; Ingólfsson, H.I.; de Jong, D.H.; Melo, M.N.; Periole, X.; de Vries, A.H.; Marrink, S.J. Dry Martini, a coarse-grained force field for lipid membrane simulations with implicit solvent. J Chem Theory Comput 2015, 11, 260–275, doi:10.1021/ct500477k.

99. Qi, Y.; Ingolfsson, H.I.; Cheng, X.; Lee, J.; Marrink, S.J.; Im, W. CHARMM-GUI Martini Maker for Coarse-Grained Simulations with the Martini Force Field. J Chem Theory Comput 2015, 11, 4486–4494, doi:10.1021/acs.jctc.5b00513.

100. Beel, A.J.; Mobley, C.K.; Kim, H.J.; Tian, F.; Hadziselimovic, A.; Jap, B.; Prestegard, J.H.; Sanders, C.R. Structural studies of the transmembrane C-terminal domain of the amyloid precursor protein (APP): does APP function as a cholesterol sensor? Biochemistry 2008, 47, 9428–9446.

101. Beel, A.J.; Sakakura, M.; Barrett, P.J.; Sanders, C.R. Direct binding of cholesterol to the amyloid precursor protein: An important interaction in lipid-Alzheimer’s disease relationships? Biochimica et biophysica acta 2010, 1801, 975–982.

102. Humphrey, W.; Dalke, A.; Schulten, K. VMD: visual molecular dynamics. Journal of molecular graphics 1996, 14, 33–38.

103. Pettersen, E.F.; Goddard, T.D.; Huang, C.C.; Couch, G.S.; Greenblatt, D.M.; Meng, E.C.; Ferrin, T.E. UCSF Chimera--a visualization system for exploratory research and analysis vesrion J Comput Chem 2004, 25, 1605–1612, doi:10.1002/jcc.20084.

