## supplemental file description for "Binding of different substrate molecules at the docking site and the active site of γ-secretase can trigger toxic events in sporadic and familial Alzheimer’s disease"

^1^ Laboratory for Biomolecular Structure and Function. Department of Biotechnology, University of Rijeka, 51000 Rijeka, Croatia.

^2^ Laboratory for Medical Biochemistry, Psychiatric Hospital Rab, Kampor 224, 51280 Rab, Croatia

**Supplement video 1. Coarse-grained molecular dynamics studies of interactions between two free C99-βCTF-APP molecules in cholesterol-lipid-bilayer.** CG structures [1] of two C99-β-CTF-APP molecules are depicted using Connolly surfaces [2]. The amino acids are colored as hydrophobic (white), positive (blue), negative (red), and polar not charged (green). The cholesterol-lipid bilayer (methods) is depicted with translucent CG spheres [1] to show cholesterol (orange) and other lipids (cyan). Yellow-cyan CG spheres represent 150 mM potassium chloride ions [1]. Multiscale MD calculations show 10 µsec of dynamic structural depictions of C99-β-CTF-APP structure that can be correlated with other studies of C99-β-CTF-APP structure [3-7]. The calculation start with fully extended C99-β-CTF-APP molecules (PDB:2LP1, [3]), i.e. no secondary structure presumption (methods). Intramolecular interactions start immediately at the start of calculations and lead to a compact structure for N-terminal and C terminal domain. The two molecules gradually show dynamic transient contacts that come and break at different sites, in a process that can be attributed to a competition between intramolecular and intermolecular interactions. The two C99-β-CTF-APP molecules ultimately form tight complementary surfaces throughout the entire protein length. Dimerization depends on the hinge point at Gly38 and Gly39 sites (green), and multiple contacts between charged and polar amino acids at the C-terminal and N-terminal domain. Notable contacts at the C-terminal domain take place between the positive Lys53-Lys54-Lys55 site and the negative site Glu74-Glu75 site. On the N-terminal domain, frequent interactions take place between Glu4 and Lys16, Arg5 and Glu22-Asp23. The transmembrane helix is hydrophobic (white) with notable polar sites at Thr 43 and Thr 48 (green). For clarity cholesterol-lipid bilayer and potassium chloride ions are shown as static spheres that have been sliced in the plane behind the protein [2]. The static dots were used to represent one out of 50 water molecules [2].

**Supplement video 2. Coarse-grained molecular dynamics studies of interactions between of free C99-β-CTF-APP substrate (PDB:2LP1, [****3****]) and γ-secretase in complex with Aβ 1-49 substrate (PDB:6IYC, [****8****]).** CG structures [1] of all proteins are depicted as partially transparent Connolly surfaces to make the interaction sites visible [2]. γ-Secretase complex shows nicastrin (pink), presenilin 1 (white), Aph1 (yellow), and Pen2 (hidden not shown). C99-βCTF-APP and Aβ 1-49 substrates are colored as hydrophobic (white), negative (red), positive (blue), and polar not charged (green). The cholesterol-lipid bilayer (methods) is depicted with translucent CG spheres [1] to show cholesterol (orange) and other lipids (cyan). Multiscale MD studies can describe 10 µsec of diffusion in the cholesterol-lipid bilayer that can be driven by the complementary electric fields (supp. Fig. 1). The corresponding changes in protein structures lead to a gradual buildup of docking interactions. The video shows specific conformations where the bound Aβ 1-49 substrate has its N-terminal maximally shielded by the nicastrin ectodomain (compare Figs. 3 and 4 in the main text). In those conditions, the closed nicastrin ectodomain can maximally block contacts between the N-terminal domains of the two substrates. Interestingly, the C-terminal domain of free C99-βCTF-APP substrate can extend towards cytosolic sections of TM6, TM6a, TM7, and the endoproteolytic site on presenilin 1. These are the most dynamic sites in presenilin structure [8,9], which control processive cleavages in Aβ production, and can be affected by the drugs and the FAD mutations that can also affect Aβ production [8-11].

**Supplement video 3. AA-MD studies of docking interactions between C-terminal section of C99-βCTF-APP substrate and cytosolic section of γ-secretase-Aβ49 complex [****8****].** The video is derived from video 2, using conversion from CG-MD to AA-MD structures [1]. The aim is to describe the docking of the C-terminal domain to the cytosolic end of presenilin to atomic details (in a molecular timeframe of 300 nanoseconds). For clarity, this video shows only C99-βCTF-APP, bound Aβ 1-49 substrate, and presenilin subunit (N-terminal gray, C-terminal pink [8]). The backbone models are used to depict protein conformers, while the transparent Connolly surfaces are used to depict protein-protein contacts [2]. The backbone of C99-βCTF-APP and bound Aβ 1-49 substrate are colored as hydrophobic (white), negative (red), positive (blue), and polar not charged (green). The red blobs represent the active sits AspH275 and Asp385 [8]. The video shows how the substrate C-terminal domain can dock with its full length to the most dynamic catalytic parts on presenilin structure, while the N-terminal domains of the two substrates is nicely separated due to the closing of nicastrin ectodomain (compare with video 2).

| 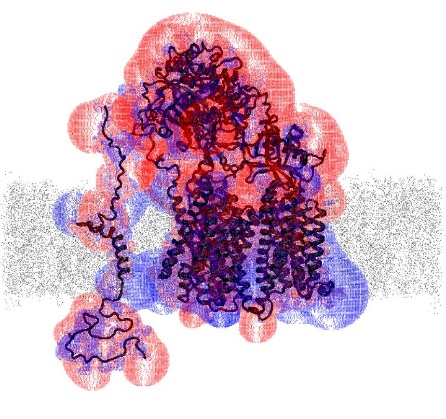 | 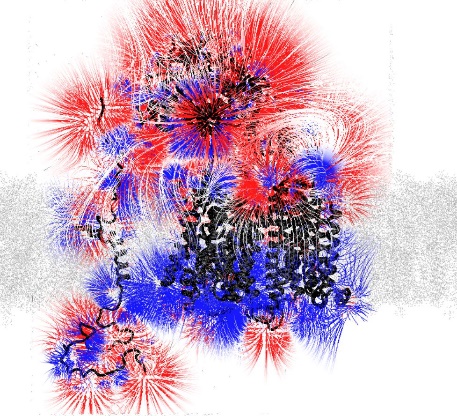 | 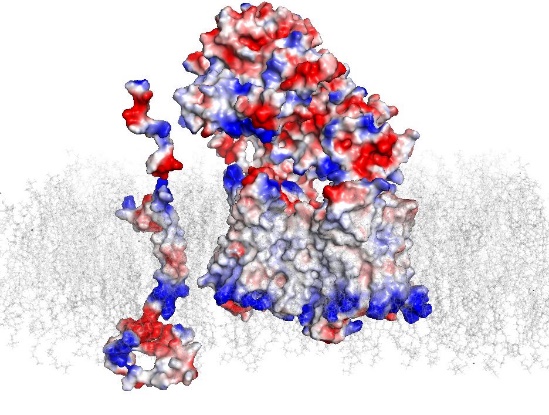 |
| --- | --- | --- |
| **A** | **B** | **C** |

**Supplement Figure 1 (A-C). Adaptive Poisson-Boltzmann Solver (APBS, [****12****]) protocols were used for the calculation of isopotential surfaces (A), field gradient lines (B), and electric potentials mapped on the Connolly surface.** Protein backbone is shown as ribbon, protein surface as Conolly surface, while silver dots show cholesterol-lipid-bilayer (methods).

(**A**) Blue and red dots represent positive and negative isopotential surfaces respectively, that are mapped in the space around the protein (scale -1.0 to 1.0 k_B_T/e).

(**B**) Blue and red lines represent gradient lines from positive to negative sites respectively, in the space around the proteins (scale -0.5 to 0.5 k_B_T/e).

(**C**) Blue and red patches represent positive and negative isopotential values respectively, that are mapped on the Connolly surface (scale -4.0 to 4.0 k_B_T/e). γ-Secretase complex is predominantly negative at the extracellular nicastrin site and predominantly positive at the intracellular presenilin at the TM6a site. The free C99-βCTF-APP substrate has complementary electric fields.

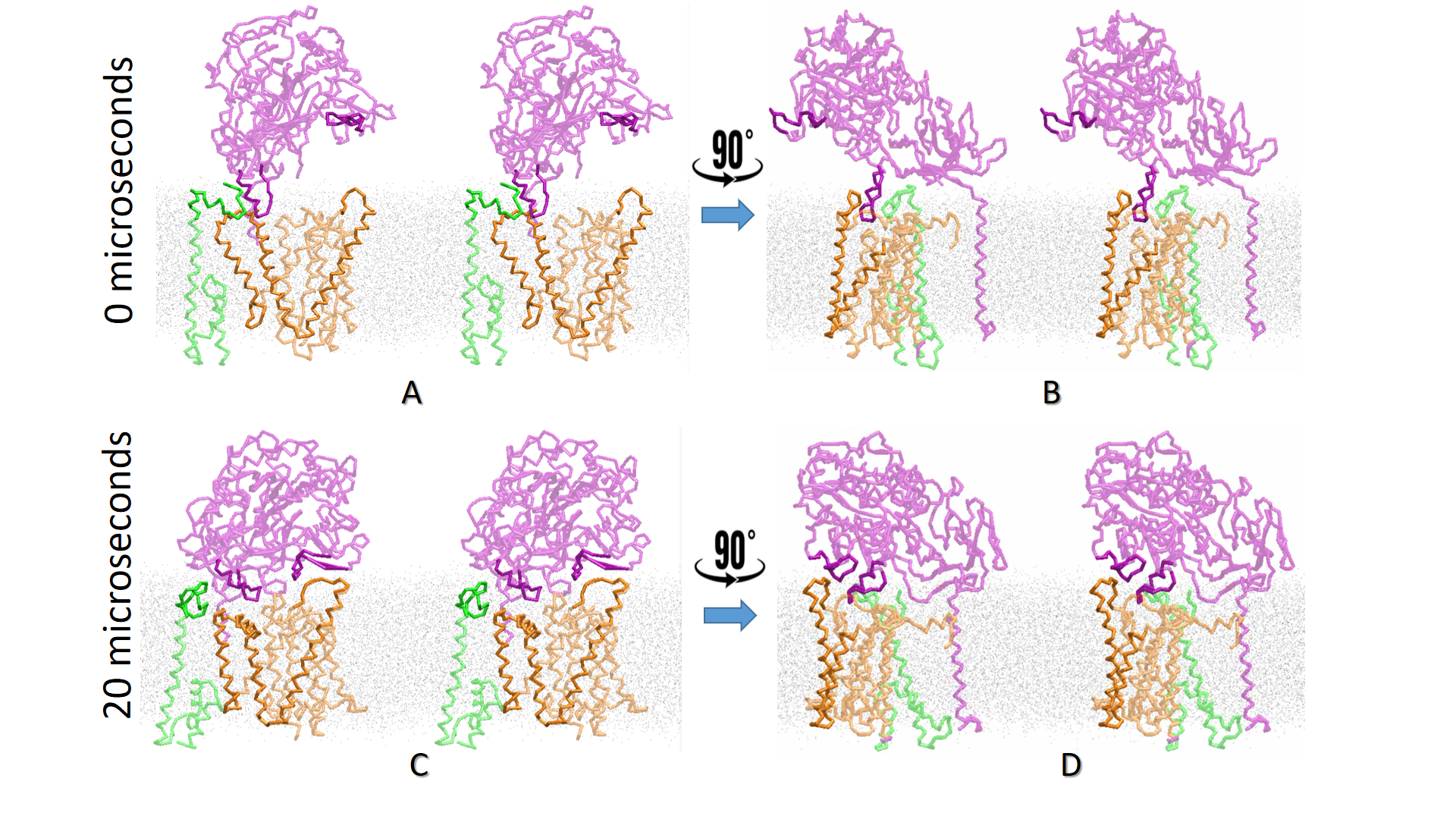

**Supplement Figure 2 (A-D). Cross-eyed stereoview view of conformational changes in γ-secretase complex upon closing of the nicastrin ectodomain.** Stereo-view is used to provide the most accurate description of conformational changes that lead to the closing of nicastrin ectodomain in γ-secretase complex. The models show nicastrin (purple), psen1 (orange), psen2 (green), and membrane (gray dots), while for clarity the Aph1 subunit is not shown. The contact points between different subunits that drive the closing of the complex are highlighted. Silver dots represents cholesterol-lipid-bilayer.

**(A-B)** EM structures show γ-secretase with the nicastrin ectodomain open ([8] PDB:6IYC).

(**C-D**) Coarse-grained molecular dynamics calculations (20 microseconds) showed that the nicastrin ectodomain can close within the first several microseconds as reported in the earlier studies [13-15]. The closing is driven by the interaction between flexible loops on nicastrin and links TM1 and TM2 on presenilin 1 (nicastrin a.a. 555-572, presenilin1 a.a. 111-123). The interaction between the two loops drives the breakdown of interaction between nicastrin and psen2 (helix a.a. 225-245 on nicastrin and a.a. 81 to 101 on presenilin 2). When psen2 is displaced, the nicastrin forms contact with TM3 (a.a. 187-195). That contacts can displace TM4 and bend the TM3 by as much as 65 degrees in the position of Phe rich sequence that starts with the Phe175. The presented interactions drive the repositioning of TM2-TM3 relative to TM4 and TM6a (C-D), and exposure of otherwise buried TM6 and TM6a (D). The closing of the nicastrin ectodomain also results in its internal twist by 27 degrees, which results in a compact structure that cannot protrude beyond the presenilin structure (compare B and D).

| 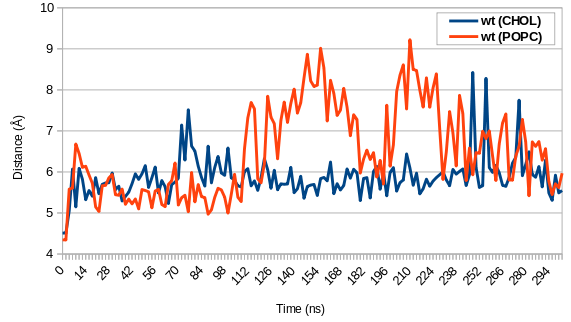 | 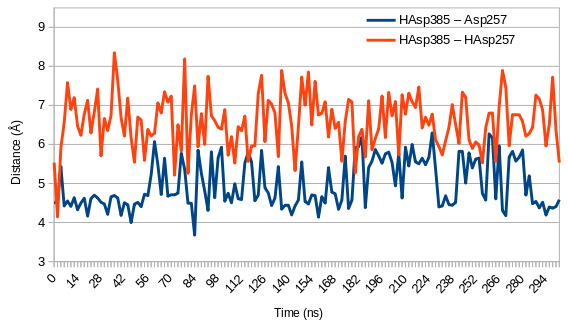 | 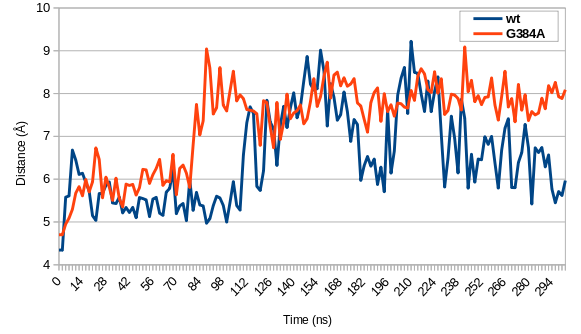 |
| --- | --- | --- |
| **A** | **B** | **C** |

**Supplement Figure 3 (A-C). AA-MD studies of active site Asp257 and Asp385 in presenilin 1.** The catalytic relevance of different MD studies can be analyzed by comparing the structure of the active site aspartates with similar studies in the past [16-18]. Namely, we measured the distance between γ-carbon on the active site aspartates [16] and we compared the calculated and the experimental pKa values [17,18].

(**A** ) The distance between γ-carbon on Asp257 and Asp385 can be affected by the lipid bilayer [19,20]. The bigger distance in POPC membranes can be attributed to higher mobility for all amino acids in proteins that are embedded in POPC bilayers (Svedružić et.al., manuscript in preparation).

(**B**) The distance between γ-carbon on the active site aspartates in cholesterol-lipid-bilayer in our studies is comparable to similar studies in the past [16]. In addition, two different protocols were used to calculate pKa values. PropKa calculations for Asp257 is 6.9 and for Asp385 is and 6.8 [17]. Delphi calculations for Asp257 is 6.9 and for Asp385 is and 6.7 [21]. These values are comparable to the experimental pKa values [18].

(**C** ) FAD mutations can affect presenilin structure, specifically the optimal catalytic distance and orientation between γ-carbon atoms on Asp257 and Asp385 (Svedružić et.al., manuscript in preparation).

References:

12.	Baker, N.A.; Sept, D.; Joseph, S.; Holst, M.J.; McCammon, J.A. Electrostatics of nanosystems: application to microtubules and the ribosome. *Proceedings of the National Academy of Sciences* **2001**, *98*, 10037-10041.

13.	Aguayo-Ortiz, R.; Chávez-García, C.; Straub, J.E.; Dominguez, L. Characterizing the structural ensemble of γ-secretase using a multiscale molecular dynamics approach. *Chem Sci* **2017**, *8*, 5576-5584, doi:10.1039/c7sc00980a.

14.	Bolduc, D.M.; Montagna, D.R.; Gu, Y.; Selkoe, D.J.; Wolfe, M.S. Nicastrin functions to sterically hinder γ-secretase-substrate interactions driven by substrate transmembrane domain. *Proceedings of the National Academy of Sciences of the United States of America* **2016**, *113*, E509-518, doi:10.1073/pnas.1512952113.

15.	Lee, J.Y.; Feng, Z.; Xie, X.Q.; Bahar, I. Allosteric Modulation of Intact γ-Secretase Structural Dynamics. *Biophys J* **2017**, *113*, 2634-2649, doi:10.1016/j.bpj.2017.10.012.

16.	Krzemińska, A.; Moliner, V.; Świderek, K. Dynamic and Electrostatic Effects on the Reaction Catalyzed by HIV-1 Protease. *Journal of the American Chemical Society* **2016**, *138*, 16283-16298, doi:10.1021/jacs.6b06856.

17.	Li, H.; Robertson, A.D.; Jensen, J.H. Very fast empirical prediction and rationalization of protein pKa values. *Proteins* **2005**, *61*, 704-721, doi:10.1002/prot.20660.

18.	Li, Y.M.; Lai, M.T.; Xu, M.; Huang, Q.; DiMuzio-Mower, J.; Sardana, M.K.; Shi, X.P.; Yin, K.C.; Shafer, J.A.; Gardell, S.J. Presenilin 1 is linked with gamma-secretase activity in the detergent solubilized state. *Proceedings of the National Academy of Sciences of the United States of America* **2000**, *97*, 6138-6143.

19.	Audagnotto, M.; Kengo Lorkowski, A.; Dal Peraro, M. Recruitment of the amyloid precursor protein by γ-secretase at the synaptic plasma membrane. *Biochem Biophys Res Commun* **2018**, *498*, 334-341, doi:10.1016/j.bbrc.2017.10.164.

20.	Aguayo-Ortiz, R.; Straub, J.E.; Dominguez, L. Influence of membrane lipid composition on the structure and activity of γ-secretase. *Physical chemistry chemical physics : PCCP* **2018**, *20*, 27294-27304, doi:10.1039/c8cp04138e.

21.	Pahari, S.; Sun, L.; Basu, S.; Alexov, E. DelPhiPKa: Including salt in the calculations and enabling polar residues to titrate. *Proteins* **2018**, *86*, 1277-1283, doi:10.1002/prot.25608.
